# The *Ptr1* locus of *Solanum lycopersicoides* confers resistance to race 1 strains of *Pseudomonas syringae* pv. tomato and to *Ralstonia pseudosolanacearum* by recognizing the type III effectors AvrRpt2/RipBN

**DOI:** 10.1101/518399

**Authors:** Carolina Mazo-Molina, Samantha Mainiero, Sara R. Hind, Christine M. Kraus, Mishi Vachev, Felicia Maviane-Macia, Magdalen Lindeberg, Surya Saha, Susan R. Strickler, Ari Feder, James J. Giovannoni, Christine D. Smart, Nemo Peeters, Gregory B. Martin

## Abstract

Race 1 strains of *Pseudomonas syringae* pv. tomato, which causes bacterial speck disease of tomato, are becoming increasingly common and no simply-inherited genetic resistance to such strains is known. We discovered that a locus in *Solanum lycopersicoides*, termed *Pseudomonas tomato race 1* (*Ptr1*), confers resistance to race 1 *Pst* strains by recognizing the type III effector AvrRpt2. In Arabidopsis, AvrRpt2 degrades the RIN4 protein thereby activating RPS2-mediated immunity. *Ptr1* also recognized homologs of AvrRpt2 from diverse bacteria including one in *Ralstonia pseudosolanacearum* and this correlated with the ability of AvrRpt2 to degrade RIN4. Using site-directed mutagenesis of AvrRpt2 we found that *Ptr1* and *RPS2* recognize identical features of AvrRpt2. However, the genome sequence of *S. lycopersicoides* revealed no *RPS2* homolog in the *Ptr1* region. *Ptr1* could play an important role in controlling bacterial speck disease and its future cloning may shed light on an example of convergent evolution for recognition of a widespread type III effector.

## Introduction

Bacterial speck disease of tomato, caused by *Pseudomonas syringae* pv. tomato *(Pst)*, occurs in cool, wet environments that favor bacterial spreading and leaf colonization through stomata (Pedley & Martin, 2003). Two races of *Pst* are currently defined which differ in their ability to cause disease on tomato varieties expressing the resistance gene *Pto*. Race 0 strains express the type III effectors AvrPto or AvrPtoB, are recognized by *Pto*, and consequently are unable to cause disease on Pto-expressing tomato varieties. Race 1 strains do not have the *avrPto* or *avrPtoB* genes or do not express these effector proteins and are therefore not recognized by *Pto* (Lin *et al*., 2006; Kunkeaw *et al*., 2010). Recently, strains with virulence attributes intermediate between race 0 and race 1 strains have been discovered (Kraus *et al*., 2017). These strains express AvrPto but nevertheless multiply to levels intermediate between race 0 and race 1 strains in tomato plants that express *Pto*.

In order to combat pathogens, plants have evolved a two-layered immune system. In an initial defense response, plants use extracellular pattern recognition receptors (PRRs) to detect the presence of microbe-associated molecular patterns (MAMPS) (Dangl *et al*., 2013). In the second immune response, plants use intracellular proteins (R proteins or nucleotide-binding oligomerization domain-like (NOD-like) receptors, NLRs) to detect pathogen effector proteins translocated inside the host cell during the infection process. NLR-triggered immunity (NTI) is typically associated with programed cell death and significant inhibition of pathogen multiplication (Jones & Dangl, 2006; Buttner, 2016).

After translocation of AvrPto or AvrPtoB into the plant cell, Pto, which encodes a serine/threonine protein kinase, physically interacts with either one of these effectors and acts in concert with Prf, an NLR, to activate NTI (Salmeron *et al*., 1996; Pedley & Martin, 2003). *Pto* was originally identified in a wild relative of tomato, *Solanum pimpinellifolium*, and it has been introgressed into many processing-type tomato varieties. For over 30 years, the gene has provided effective control of speck disease caused by *Pst* race 0 strains (Pitblado & Kerr, 1980). However, the increasing prevalence of race 1 *Pst* strains able to overcome Pto/Prf-mediated NTI has led to the search for sources of resistance to these virulent strains. Since no race 1 resistance has been identified in cultivated tomato, wild relatives of tomato are likely to be the best potential source for this trait (Peralta *et al*., 2008).

Wild relatives of tomato have been screened previously to identify resistance against race 1 *Pst* strains. One study reported a screen of introgression lines derived from *S. habrochaites* LA1777 using the race 1 strain A9 from California (Thapa *et al*., 2015). The detection of four QTLs, on chromosomes 1, 2 and 12 (2 loci), explained the moderate resistance to this strain, however, overall they accounted for a small percentage of the variability observed (10.5–12.5 % of the phenotypic variation) (Thapa *et al*., 2015). A second study identified two QTLs, on chromosomes 2 and 8, in *S. habrochaites* accession LA2109 that contributed to resistance to race 1 strain T1, which accounted for 24% and 26% of the phenotypic variability, respectively (Bao *et al*., 2015). Recently, another study reported a screen of 96 wild accessions and identified two accessions that display resistance toward race 1 strain T1, *S. neorickii* LA1329 and *S. habrochaites* LA1253. Resistance in LA1253 appears to be a complex genetic trait and its inheritance remains unclear (Hassan *et al*., 2017). Although together these QTLs might contribute to the breeding of enhanced race 1 *Pst* resistance in tomato, their quantitative nature and relatively weak race 1 resistance limits their usefulness.

Here we report the identification of race 1 resistance in a distantly-related relative of tomato, *Solanum lycopersicoides*. The resistance is likely due to a single locus which we refer to as *<u>P</u>seudomonas <u>t</u>omato <u>r</u>ace <u>1</u>* (*Ptr1*). *Ptr1* confers resistance to several race 1 *Pst* strains but not to the race 0 strain DC3000. A test of type III effectors that are present in race 1 strains but lacking in DC3000 identified AvrRpt2 as the effector recognized by *Ptr1*. AvrRpt2 is a cysteine protease that has been intensively studied because it cleaves RIN4 leading to activation of the Arabidopsis NLR protein RPS2 (Axtell *et al*., 2003; Mackey *et al*., 2003). By using site-directed mutagenesis of AvrRpt2 we discovered that *Ptr1* and *RPS2* recognize identical features of this effector. *Ptr1* also recognizes AvrRpt2 variants expressed by diverse bacterial phytopathogens and this recognition correlates with ability of the variants to cleave the Arabidopsis RIN4 protein. Notably, Ptr1 conferred strong resistance to a *Ralstonia pseudosolanacearum* strain that expresses an AvrRpt2 homolog (RipBN); no resistance gene that gives protection against this important pathogen, causative agent of bacterial wilt disease, has been reported previously.

## Materials and Methods

### Bacterial strains and plasmids

*Pseudomonas syringae* pv. tomato strains DC3000 (Buell *et al*., 2003), NY15125 (Kraus *et al*., 2017), T1 (Almeida *et al*., 2009), JL1065 (Whalen *et al*., 1991), NYT1 (Jones *et al*., 2015), CA-A9, and CA-407 (Kunkeaw *et al*., 2010) were grown on King’s B (KB) (King *et al*., 1954) semi-selective media at 30°C (Table S1). *Ralstonia pseudosolanacearum* CMR15 (Remenant *et al*., 2010) was grown on rich B medium (10 g l^−1^ bacto-peptone, 1 g l^−1^ yeast extract, 1 g l^−1^ casamino acids). Plasmids pCPP5372 (Oh *et al*., 2007) carrying wildtype *avrRpt2NY15125, avrRpt2* variants or the empty vector were introduced into DC3000 by electroporation (Table S2 and S3). All *P. syringae* pv. tomato strains were stored in 20% glycerol + 60 mM sucrose at – 80°C. *Escherichia coli* TOP10 was used for plasmid maintenance and grown in LB medium at 37°C.

### Plant material

*Solanum lycopersicoides* introgression lines seeds of LA4245 and LA4277 were obtained from the Tomato Genetics Resource Center (https://tgrc.ucdavis.edu/lycopersicoides_ils.aspx). The genotype of LA4245 can be heterozygous for the presence of *Ptr1 (Ptr1/ptr1)* (LA4245-R) or homozygous for the lack of the gene *(ptr1/ptr1*) (LA4245-S). *S. lycopersicoides* introgression lines were grown in a greenhouse at 24°C during daylight and 22°C at night. *Nicotiana benthamiana* Nb-1 (Bombarely *et al*., 2012) was maintained in a growth chamber with 16 h : 8h, light : dark at 24°C with light and 20°C in the dark and 50% humidity. Tomatoes and *N. benthamiana* plants were grown in Cornell Osmocote Mix soil (0.16 m^3^ peat moss, 0.34 m^3^ vermiculite, 2.27 kg lime, 2.27 kg Osmocote Plus15-9-12 and 0.54 kg Uni-Mix 11-5-11; Everris, Israeli Chemicals Ltd). After pathogen inoculation, plants were moved to a growth chamber with 25°C, 50% humidity, and 16 h light. *Arabidopsis thaliana* accession Columbia (Col-0) seeds were ethanol-sterilized, suspended in 1 ml of water and cold stratified for 2 days at 4°C. *A. thaliana* was grown in Fafard Mix (Sungro Horticulture) in a growth chamber under fluorescent lighting (100 μmol m^−2^ s^−1^) with a 12 h : 12 h; light : dark cycle at 21°C and 40% humidity.

### Genome sequencing, assembly, and type III effector annotation of NY15125

The NY15125 genome was sequenced to 163X coverage with long reads from the Pacbio RSII platform. A Canu assembly was performed with a stringent error rate (correctedErrorRate = 0.035) (Koren *et al*., 2017). Illumina sequencing was also done to generate paired-end reads for a coverage of 114X. Adapter clipping and quality filtering of the Illumina reads was done with Trimmomatic (Bolger *et al*., 2014). Concordant read mapping of the Illumina paired-end reads was used to evaluate the quality of the assembly. Two rounds of base level error corrections were done with Pacbio reads using Arrow (https://github.com/PacificBiosciences/pbbioconda), followed by two rounds of error correction with Illumina reads using Pilon (Walker *et al*., 2014). The polished assembly includes a 6.2 Mb chromosome and three putative plasmids (88Kb, 116Kb, 122Kb). It was annotated with Prokka (Seemann, 2014) using proteins from the T1 genome (Almeida *et al*., 2009) as supporting evidence. The NY15125 chromosome was compared with the DC3000 and T1 genomes using BRIGG (Alikhan *et al*., 2011). Pseudomolecules were constructed from the deposited contigs for *P. syringae* pv. tomato NY15125 and annotated using MG-RAST (Meyer *et al*., 2008). Effector genes were identified from the MG-RAST annotation, by alignment with other *P. syringae* pv. tomato sequences, and based on proximity to HrpL binding sites, predicted using the methods described previously (Saha & Lindeberg, 2013). The NY15125 genome sequence and plasmid sequences are available from GenBank (accession numbers: CP034558-CP034561).

### Development of a *P. syringae* pv. tomato NY15125Δ*avr*Rpt2 strain

A 1,024-bp promoter fragment and a 841-bp fragment downstream of the *avrRpt2* gene sequence were PCR amplified and *EcoRI* or *XmaI* restriction sites were added, respectively. Fusion of both DNA fragments was cloned into the suicide vector pK18mobsacB and transformed into *E. coli* S17-1. Deletion of *avrRpt2* in NY15125 was performed by biparental mating as described previously (Kraus *et al*., 2017) with modifications (Kvitko & Collmer, 2011).

### *P. syringae* pv. tomato inoculation and population assays in tomato

*P. syringae* pv. tomato strains were grown on KB plates for 2 days at 30°C. Strains were diluted in 10 mM MgCl_2_ + 0.002% Silwet L-77 at a final concentration of 5 x 10^4^ cfu ml^−1^. Four-week-old plants were vacuum infiltrated, and three leaf disk samples (7 mm in diameter) were collected at 2 h (day 0) and 2 days post inoculation (dpi) to quantify bacterial populations. The experiments were repeated three times. Results shown are the mean of three independent experiments using three biological replicates per strain, including standard error of the mean. Photographs for each technical replicate were taken 7 dpi. Statistical analyses were performed using Prism 6.0 (GraphPad Software).

### *P. syringae* pv. tomato inoculation and population assays in *Arabidopsis thaliana*

Five-week-old plants were dip inoculated for 20 seconds in a bacterial suspension (1 x 10^8^ cfu ml’^1^ of *Pst*) containing 10 mM MgCl_2_ + 0.02% Silwet L-77. Bacterial populations were measured 3 dpi by submerging the aerial plant tissue in 10 mM MgCl_2_ + 0.2% Silwet L-77 for 2 h at 28°C. The bathing solution was serially diluted and plated (Tornero & Dangl, 2001).The experiments were repeated three times. Results shown are the mean of three independent experiments using three biological replicates per strain, including standard error of the mean. Statistical analyses were performed using Prism 6.0 (GraphPad Software).

### Immunodetection of AvrRpt2 proteins in *P. syringae* pv. tomato

Strains of *P. syringae* pv. tomato grown on KB plates for 2 days at 30 °C were resuspended in hrp-inducing liquid minimal media (50 mM KH_2_PO_4_, 1 g l^−1^ (NH_4_)_2_SO_4_, 1.7 mM NaCl, 1.7 mM MgCl_2_, pH 5.7) or KB liquid media containing the appropriate antibiotics at an OD_600_ of 0.4 and 0.1 respectively. Bacterial cultures were grown at 28°C for 16 hours shaking at 220 RPM and OD_600_ was adjusted to a final concentration of 0.5. One ml of each bacterial culture was washed with water and centrifuged. Bacterial pellets were resuspended in 100 μl of Laemmli buffer (20 mM Tris, 1% sodium dodecyl sulfate [SDS], 0.05% bromphenol blue, and 10% glycerol, pH 6.8), boiled for 5 minutes, and 5 μl of each was used for immunoblot analysis. To detect AvrRpt2 proteins, membranes were probed with ∝-HA antibody (Roche, Indianapolis, IN) conjugated with HRP.

### Agrobacterium-mediated transient protein expression in leaves

*Agrobacterium tumefaciens* 1D1249 (Wroblewski *et al*., 2005) harboring the various expression vectors were grown on LB media with the appropriate antibiotics for 2 days at 30°C. Bacteria were scraped from the plate, resuspended in infiltration buffer (10 mM MgCl2, 10 mM MES [pH 5.6], and 200 mM acetosyringone) and maintained for 4 h in the dark at room temperature on a nutator rocker. Bacterial cultures were then washed, centrifuged, and the pellet was resuspended in fresh infiltration buffer before diluting cultures at a final OD_600_ of 0.15. Tomato or *N. benthamiana* leaves were infiltrated using a needle-less syringe and placed to a growth chamber (24°C day and 22°C night). Leaf samples for protein expression were taken 32 hours later.

### Immunoblot detection of plant-expressed proteins

Protein samples were analyzed by grinding three leaf disks (9 mm in diameter) in protein sample buffer (50 mM Tris HCl [pH 7.5], 10% glycerol, 2% SDS, 2 mM EDTA, 1 mM DTT and 1% protease inhibitor [Sigma-Aldrich]). Samples were separated by SDS-PAGE on 4-20% gradient polyacrylamide gels and transferred to Immobilon-P PVDF membranes (Millipore) according to standard procedures (Taylor, 2015). To detect AvrRpt2 proteins, membranes were probed with ∝-c-Myc (GeneScript, Piscataway, NJ) antibody conjugated with HRP. For tomato Rin4 detection, *At*Rin4 polyclonal antiserum (gift from G. Coaker, UC-Davis) was used at a concentration of 1:2.000. Secondary goat anti-rabbit IgG conjugated with HRP was used at a dilution of 1:10,000 (Promega, Madison, WI).

### *Ralstonia pseudosolanacearum* disease assays

For survival assays with *R. pseudosolanacearum* CMR15, 4-week-old tomato plants (grown in peat pots) were transferred in potting mixture in 3-liter pots to the Toulouse Plant-Microbe Phenotyping facility, (28°C, 16 hours light). Each tomato plant was soil-drench inoculated with 50 ml of 10^8^ cfu ml^−1^. Disease scoring was performed daily using a visual index in which the numbers 1, 2, 3 and 4 corresponded to 25%, 50%, 75% and 100% wilted leaves, respectively. Disease scores were transformed into binary data for the purpose of statistical comparison between disease curves (Remigi *et al*., 2011).

### Tomato genome sequencing

Genomic DNA was extracted from a single LA4245-R plant using a DNeasy Plant Mini Kit (QIAGEN). DNA was mechanically sheared using the Covaris S2 Adaptative Focused Acoustic Disruptor (Covaris, Inc., Woburn, MA, USA) to an average size of 500-600 bp, and used to prepare a library. Single-end 100 bp DNA reads were sequenced using the Illumina HiSeq 2000 platform. The reads from LA4245-R were mapped to the *S. lycopersicum* Heinz 1706 genome sequence SL2.50 (Tomato Genome Consortium, 2012) using hisat2 version 2.1.0 (Kim *et al*., 2015), and SNPs were called using GATK version 4.0. (McKenna *et al*., 2010). SNPs were plotted in 10 kb bins using R. Synteny between SL2.50 and LA2951 version 0.6 was determined using SynMap at CoGe: https://genomevolution.org/coge/SynMap.pl. Protein similarity for RPS2 and MR5 were calculated with Geneious R11: https://www.geneious.com. The genome sequence of LA4245 is available at NCBI SRA under project ID xxxx. The genome sequence of LA2951 is available at: https://solgenomics.net/organism/Solanum_lycopersicoides/genome.

## Results

### A locus on chromosome 4 from *S. lycopersicoides* LA2951 confers resistance to *Pst* race 1 strains

During the summer of 2015 in upstate New York, a research plot of 110 *S. lycopersicum* VF36 x *S. lycopersicoides* LA2951 introgression lines (ILs; (Canady *et al*., 2005)) became naturally infected by *Pseudomonas syringae* pv. tomato (*Pst*) resulting in severe symptoms of bacterial speck disease. However, two ILs, LA4245 and LA4277, remained essentially free of disease. LA4245 and LA4277 have large overlapping introgressed segments from chromosome 4 of *S. lycopersicoides* (Canady *et al*., 2005). The introgression in LA4245 is smaller and so we focused on that line for further characterization. In order to determine the race and other characteristics of the *Pst* strain involved in the outbreak, isolates were collected from the field and analyzed. The presence of both *avrPto* and *avrPtoB* genes, immunoblot detection of the AvrPto protein, and subsequent inoculation of tomato plants with and without the *Pto* gene indicated the field isolates were race 0 *Pst* strains (Kraus *et al*., 2017). One *Pst* strain, referred to as NY15125, was chosen for further analysis.

The chromosome 4 introgression segment in LA4245 is maintained in heterozygous condition because homozygotes are very rarely obtained, as noted previously (Canady *et al*., 2005). We named the putative LA4245 resistance locus *Ptr1* (*<u>P</u>seudomonas <u>tomato</u> <u>r</u>ace 1*) and used a nomenclature in which its presence or absence is denoted as LA4245-R (*Ptr1/ptr1*) or LA4245-S (*ptr1/ptr1*), respectively. To follow up the field observations, LA4245 plants were inoculated with *Pst* strains DC3000, NY15125 and T1 in the greenhouse. Pathogen assays showed that DC3000 caused severe symptoms on LA4245-R plants, whereas NY15125 and T1 caused the appearance of few or no specks on LA4245-R, respectively (Fig. **1a**). All of the *Pst* strains caused more disease on LA4245-S plants. Consistent with these observations, bacterial population assays showed that T1 attained levels ~65-fold lower than DC3000 on LA4245-R plants (Fig. **1b**). NY15125 grew to a level intermediate between T1 and DC3000 on LA4245-R. Such intermediate growth was observed previously for NY15125 on Pto-expressing plants (Kraus *et al*., 2017). The three *Pst* strains all attained similar population levels on LA4245-S plants (Fig. **1b**). Therefore, the putative *Ptr1* locus from *S. lycopersicoides* LA2951 confers resistance to *Pst* strains NY15125 and notably T1, which is a race 1 strain.

**Fig. 1.**
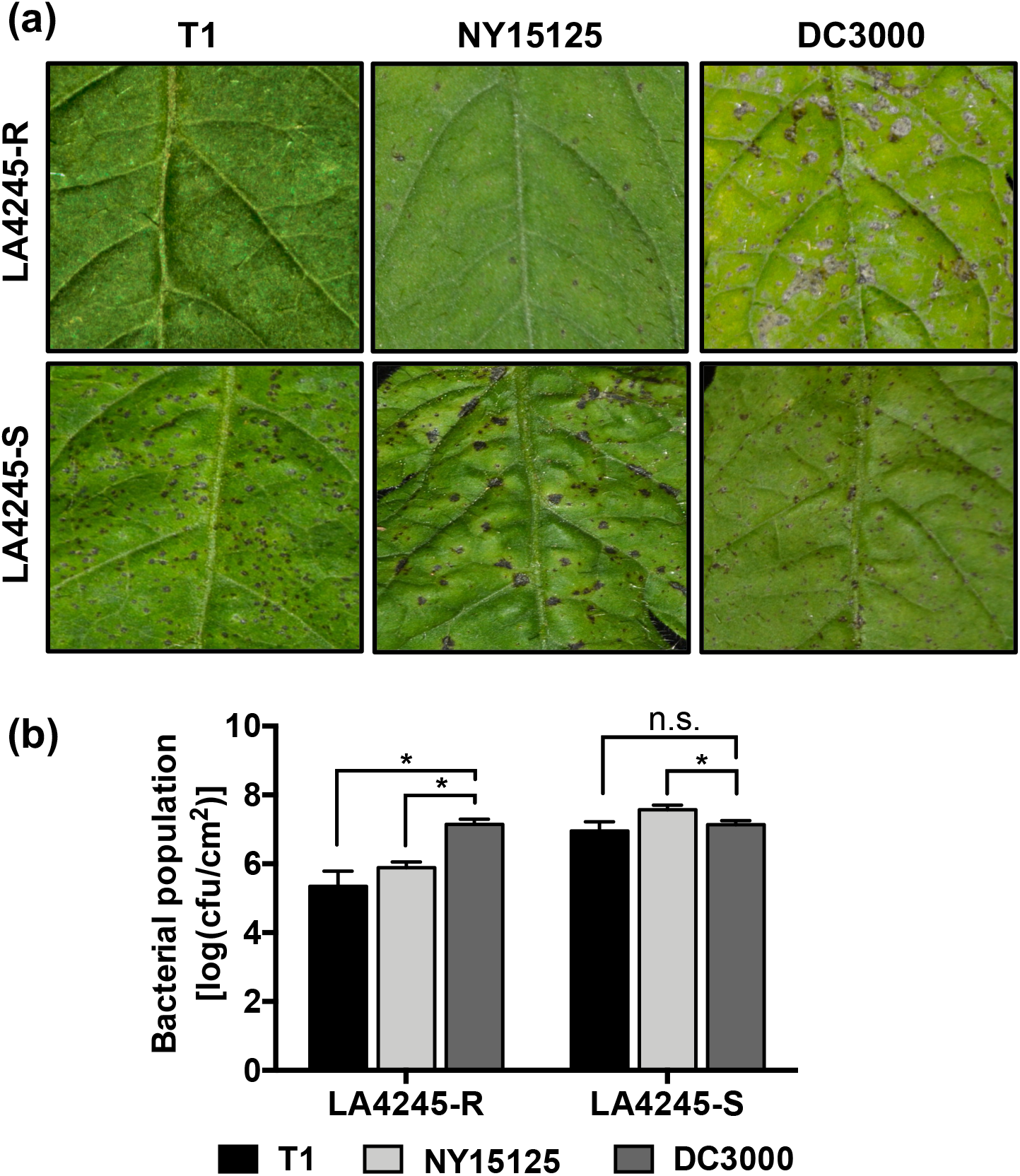
LA4245-R confers resistance to certain strains of *Pseudomonas syringae* pv. tomato. (a) Disease symptoms of LA4245-R and LA4245-S leaves vacuum infiltrated with DC3000, NY15125 and T1 at 3 x 10^4^ cfu ml^−1^and photographed five days later. (b) Bacterial populations measured two days after inoculation with the *Pst* strains indicated. Significance was determined by a pair-wise t-test and is indicated as: *significant at *P*<0.05, not significant (n.s.) at *P*>0.05. Bars indicate the mean of three independent experiments using three plants per strain. Error bars represent +/− SEM.

### *Pst* resistance in LA4245-R plants is due to recognition of the bacterial effector AvrRpt2

A comparison of the type III effector genes in DC3000 and T1 identified eight that are present exclusively in T1 (*avrA1, avrRpt2, hopAE1, hopAG1, hopAI1, hopAS1, hopS1*, and *hopW1*) (Jones *et al*., 2015). To determine whether LA4245-R resistance involves the recognition of any of these effectors, T1-specific effectors were individually cloned into the expression vector pCPP5372, the plasmids were introduced via electroporation into DC3000 Δ*avrPto*Δ*avrPtoB* and the strains were inoculated onto LA4245-R plants. All of the strains, except the one expressing AvrRpt2, caused disease on LA4245-R plants. Subsequent experiments showed that the DC3000 strain expressing AvrRpt2 reached a population size 80-fold less in leaves of LA4245-R plants compared to LA4245-S plants; a DC3000 strain carrying an empty vector grew to the same level in the two plant lines (Fig. **2a**).

**Fig. 2.**
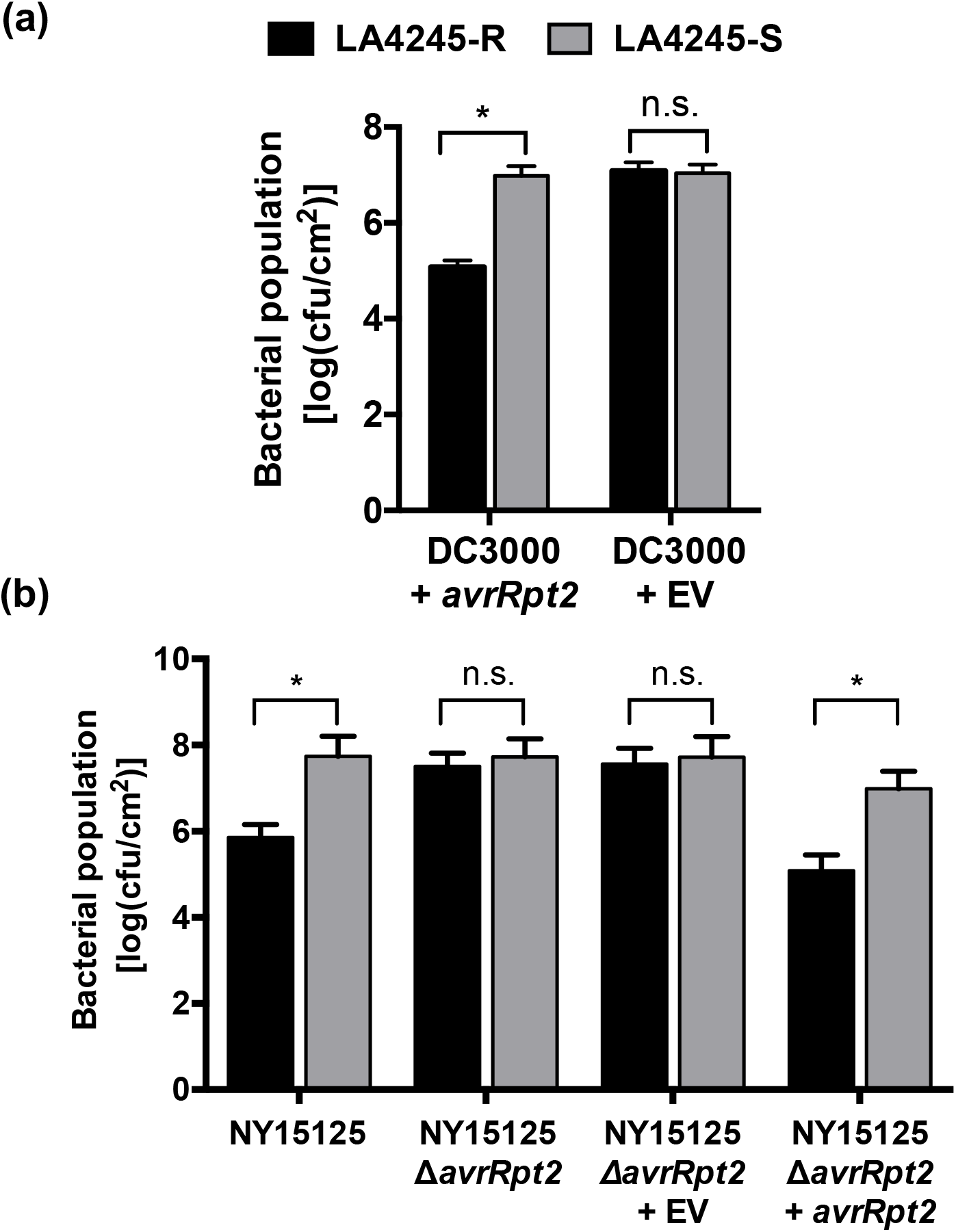
*Pst* race 1 resistance in LA4245-R plants is due to the recognition of the bacterial effector AvrRpt2. (a) LA4245-R and LA4245-S plants were vacuum infiltrated with DC3000 carrying *avrRpt2NY15125* or an empty vector (EV) at 5 x 10^4^ cfu ml^-1^and bacterial populations were measured two days later. (b) LA4245-R and LA4245-S plants inoculated with NY15125 wild-type, NY15125Δ*avrRpt2*, a complemented strain, or an empty vector (EV) at 5 x 10^4^ cfu ml^−1^. Bacterial populations were measured two days after inoculation. Significance was determined by a pair-wise t-test and is indicated as: * significant at *P*<0.05, not significant (n.s.) at *P*>0.05. Both (a) and (b) bars indicate the mean of three independent experiments using three plants per strain. Error bars represent +/− SEM.

Next, we asked whether *Pst* isolate NY15125, which was collected from the naturally-infected tomato field, has the *avrRpt2* gene. Genome sequencing and PCR analysis confirmed the presence of *avrRpt2* in this isolate (Fig. S1, Table S4). A comparison of AvrRpt2 protein sequences from NY15125 and T1 showed they are 100% identical. To investigate the activity of *avrRpt2* in NY15125, we generated a NY15125*DavrRpt2* deletion mutant and tested whether the strain was altered in its ability to grow on LA4245-R or LA4245-S plants. LA4245-R plants inoculated with the NY15125 wild-type strain supported a ~60-fold lower bacterial population compared to LA4245-S plants (Fig. **2b**). In contrast, this difference was not observed when LA4245-R and LA4245-S plants were inoculated with NY15125*DavrRpt2*. Importantly, complementation of the mutant strain with *avrRpt2* restored the wild-type growth difference in LA4245-R plants (Fig. **2b**).

AvrRpt2 was originally identified in *Pst* JL1065 (Whalen *et al*., 1991). A comparison between the AvrRpt2 protein sequences from JL1065 and NY15125 showed that the proteins have just two divergent amino acids: proline-24 and alanine-152 in AvrRpt2_JL1065_ are replaced by threonine and glycine, respectively in AvrRpt2_NY15125_. We therefore investigated if there was any difference in effector recognition on LA4245-R plants when infected with JL1065 wild-type and JL1065Δ*avrRpt2* strains (Lim & Kunkel, 2005). Bacterial population assays indicated that AvrRpt2_JL1065_ is also recognized by LA4245-R (Fig. S2). Finally, we tested three additional race 1 *Pst* strains which all carry the *avrRpt2* gene (NYT1, CA-A9, and CA-407) and found that LA4245-R is resistant to all of them, whereas LA4245-S was susceptible to these strains (Fig. S3). Thus, *Ptr1* confers AvrRpt2-mediated resistance to multiple *Pst* race 1 strains.

### The *Ptr1* locus and *RPS2* recognize identical features of AvrRpt2

To gain insight into the mechanism of *Ptr1* recognition of AvrRpt2, we performed site-directed mutagenesis to alter amino acids in the effector that have been reported to be essential for its recognition by RPS2 in Arabidopsis (Jin *et al*., 2003; Lim & Kunkel, 2004a; Lim & Kunkel, 2004b; Chisholm *et al*., 2005). Ten AvrRpt2 variants were generated, of which eight have been reported to abolish recognition by RPS2 (Axtell *et al*., 2001; Axtell *et al*., 2003; Jin *et al*., 2003; Lim & Kunkel, 2004a); two variants, F70R, which disrupts the AvrRpt2 autocleavage site, and E150S are still recognized by RPS2 (Jin *et al*., 2003; Chisholm *et al*., 2005). Each AvrRpt2 variant was introduced into DC3000 on a plasmid and shown to be expressed by immunoblotting (Fig. S4). The strains were then vacuum infiltrated into LA4245-R and LA4245-S plants and bacterial populations in leaves were measured and plants scored for disease symptoms. We observed that DC3000 expressing AvrRpt2 variants with the substitutions C122A, C122Y, G131D, G141D, G194R or H208A grew to high levels and caused severe disease on LA4245-R plants, indicating a loss of *Ptr1* recognition (Fig. 3**a,b**). In contrast, similar to wild-type AvrRpt2, strains with variants F70R, E150S, Y191C or D216E reached population levels on average ~60-fold lower than the empty vector control strain and did not cause any disease symptoms on LA4245-R plants (Fig. 3**a,b**). All of the DC3000 strains caused disease and reached similar population levels in LA4245-S plants (Fig. S5).

**Fig. 3.**
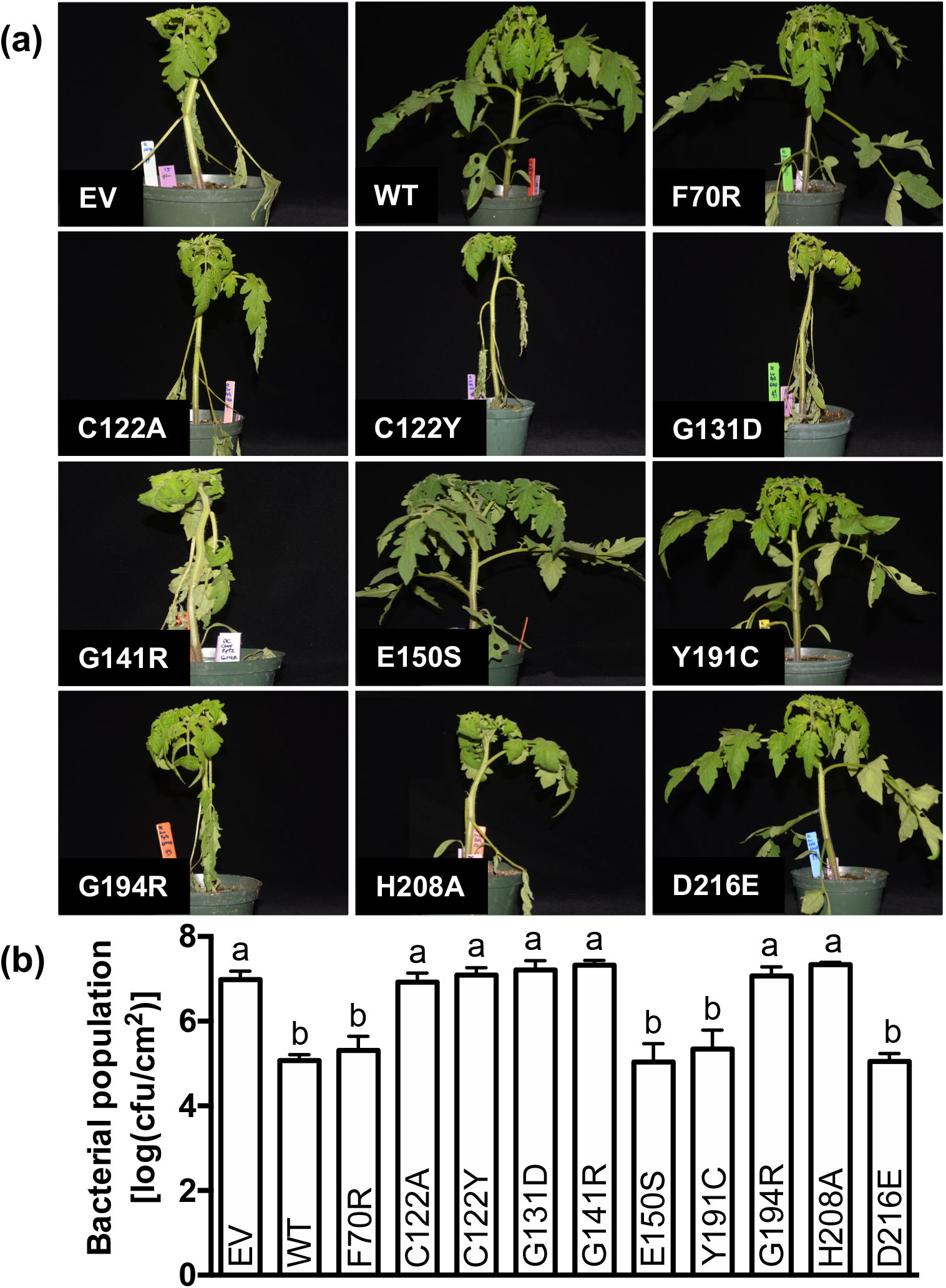
*Ptr1* detects the same features of AvrRpt2 as does *RPS2*. (a) Symptoms of LA4245-R plants vacuum infiltrated with DC3000 expressing *avrRpt2* wild-type (WT), *avrRpt2* variants or an empty vector (EV) at 5 x 10^4^ cfu ml^−1^. Photographs were taken five days after inoculation. (b) LA4245-R plants inoculated with DC3000 expressing *avrRpt2* wild-type (WT), *avrRpt2* variants and empty vector (EV). Bacterial population were measured two days after inoculation. Significance was determined using ANOVA with a Tukey’s post hoc multiple comparison test, and different letters indicate significant differences between treatments (*P*<0.001). Bars indicate the mean of three independent experiments using three plants per strain. Error bars represent +/− SEM.

Interestingly, AvrRpt2 variants with the substitutions Y191C and D216E were recognized by *Ptr1* but have been reported earlier not to be recognized by *RPS2* and to be unable to induce Arabidopsis RIN4 degradation (Lim & Kunkel, 2004b). However, recently it was shown that Arabidopsis RIN4 is cleaved by variants AvrRpt2(Y191C) and AvrRpt2(D216E) (Eschen-Lippold *et al*., 2016). In order to understand this discrepancy, Arabidopsis Col-0 *RPS2* plants were inoculated with DC3000 expressing AvrRpt2 and several of the variants, including Y191C and D216E, and bacterial population were measured three days later. We observed a significant reduction in bacterial growth and an absence of disease symptoms in Col-0 *RPS2* plants inoculated with DC3000 carrying AvrRpt2 and the variants E150S, Y191C, and D216E; the F70R variant appeared to be weakly detected by RPS2 (Fig. S6). Thus *Ptr1* and *RPS2* detect the same features of AvrRpt2.

### *Ptr1* recognition of the AvrRpt2 variants correlates with the effector’s ability to cleave tomato Rin4 proteins

In Arabidopsis, AvrRpt2-mediated degradation of RIN4 leads to the activation of RPS2 (Mackey *et al*., 2003). Accordingly, we hypothesized that AvrRpt2 variants that are recognized by *Ptr1* will also be capable of degrading Rin4 in tomato. Tomato has three genes with similarity to *AtRIN4* that are expressed in leaves and two of these are induced during Pto-mediated NTI (*SlRin4-1, SlRin4-2*, and *SlRin4-3*; Table S5). To avoid the Ptr1-mediated defense response, we vacuum infiltrated LA4245-S plants with the DC3000 strains expressing each AvrRpt2 variant and wildtype AvrRpt2 and detected SlRin4 abundance by immunoblotting 6 hours later. Wild-type AvrRpt2 and the variants F70R, E150S, Y191S and, D216A each induced a reduction in tomato Rin4 abundance which correlates with *Ptr1* recognition of these proteins in LA4245-R plants (Fig. **4a**). DC3000 strains with AvrRpt2 variants C122A, G131D, G141D, G194R, H208A, or an empty vector, all of which caused disease on LA4245-R plants, failed to induce tomato Rin4 elimination (Fig. **4a**). Similar experiments in Arabidopsis, supported our earlier observations in that both AvrRpt2(Y191C) and AvrRpt2(D216E) were able to degrade RIN4 in *rps2* plants (Fig. **4b**). Therefore, there is a perfect correlation between the recognition of each AvrRpt2 variant by *Ptr1* and its ability to induce tomato Rin4 disappearance.

**Fig. 4.**
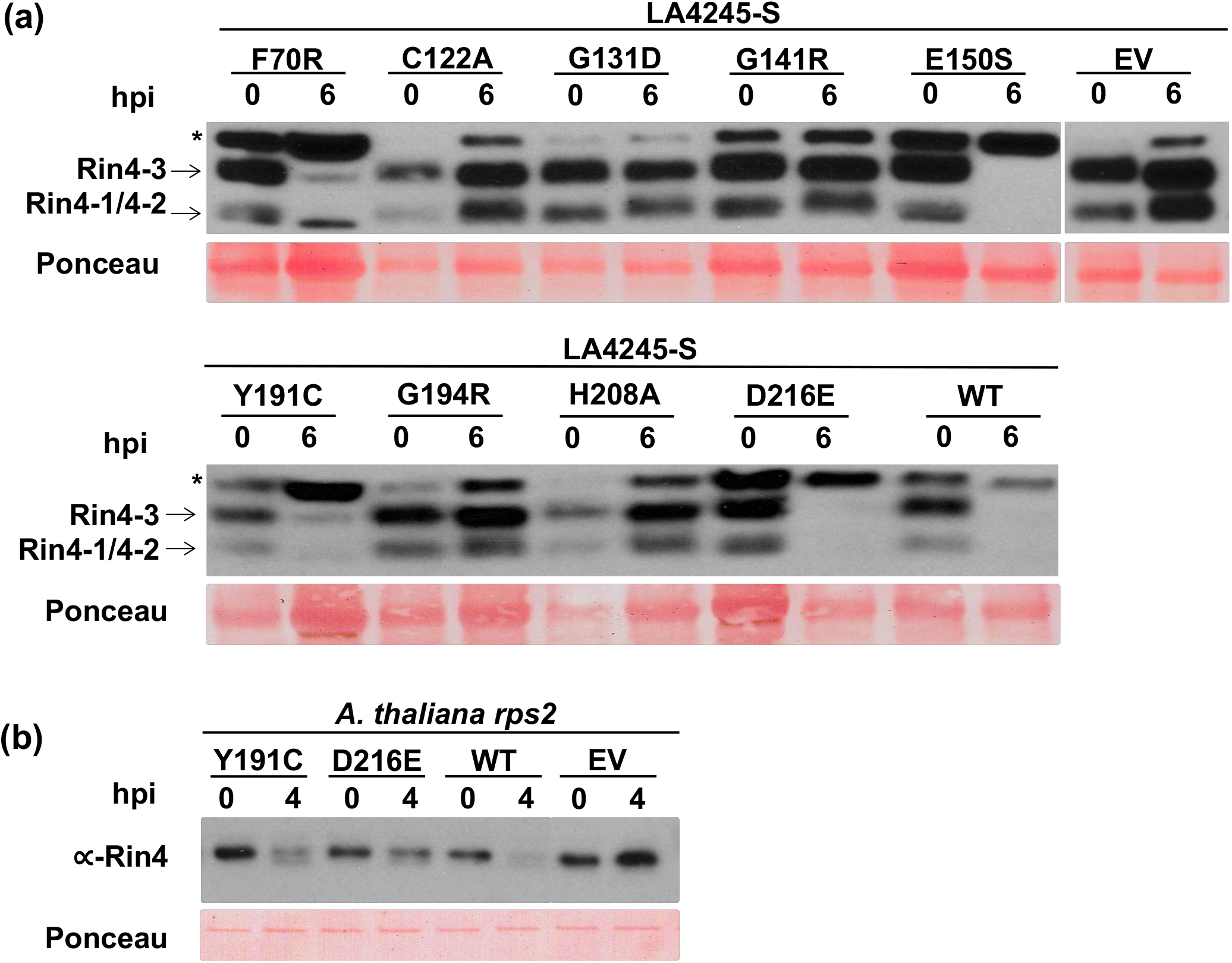
AvrRpt2 variants F70R, E150S, Y191C, D216E and AvrRpt2 wild-type which are recognized by *Ptr1* are able to degrade tomato Rin4 proteins. (a) Degradation of the endogenous tomato Rin4 proteins after vacuum infiltration of LA4245-S leaves with DC3000 strains carrying *avrRpt2* wild-type (WT), *avrRpt2* variants or an empty vector (EV). Plant tissue was harvested 0 and 6 hours post infiltration (hpi). Total protein extracted from infiltrated plants was subjected to immunoblotting using an ∝-RIN4 antibody. Locations of tomato Rin4-1, Rin4-2, and Rin4-3 are shown. The asterisk indicates an unknown cross-reacting protein. (b) Arabidopsis Col-0 *rps2* leaves were syringe-infiltrated with DC3000 carrying *avrRpt2* wild-type (WT), *avrRpt2* variants or an empty vector (EV). Leaf tissue was harvested 4 hours after infiltration. Total protein extracted from infiltrated leaves was subjected to immunoblotting using an ∝-RIN4 antibody. Ponceau staining shows amount of protein loaded in each lane.

### AvrRpt2 proteins from different bacterial species are recognized by *Ptr1*

Homologs of AvrRpt2 are found in diverse bacterial species including the plant pathogens *Erwinia amylovora, Ralstonia pseudosolanacearum, Acidovorax citrulli* and *Acidovorax avenae*, the soil bacterium *Burkholderia pyrrocinia*, the fungal parasite *Collimonas fungivorans*, and the symbiotic bacteria *Mezorhizobium huakuii* and *Sinorhizobium medicae* (Zhao *et al*., 2006; Eschen-Lippold *et al*., 2016). Some of these AvrRpt2 proteins have very divergent amino acid sequences and we wished to determine whether *Ptr1* would recognize them. *Agrobacterium-mediated* expression (agroinfiltration) was used to express each AvrRpt2 protein or a YFP control in leaves of LA4245-R and LA4245-S. The AvrRpt2 homologs from five of the eight bacterial species induced cell death in LA4245-R leaves but not in LA4245-S leaves indicating they are recognized by *Ptr1* (Fig. **5**). Detection of protein expression of each AvrRpt2 homolog was done via agroinfiltration in *N. benthamiana* leaves (Fig. S7).

**Fig. 5.**
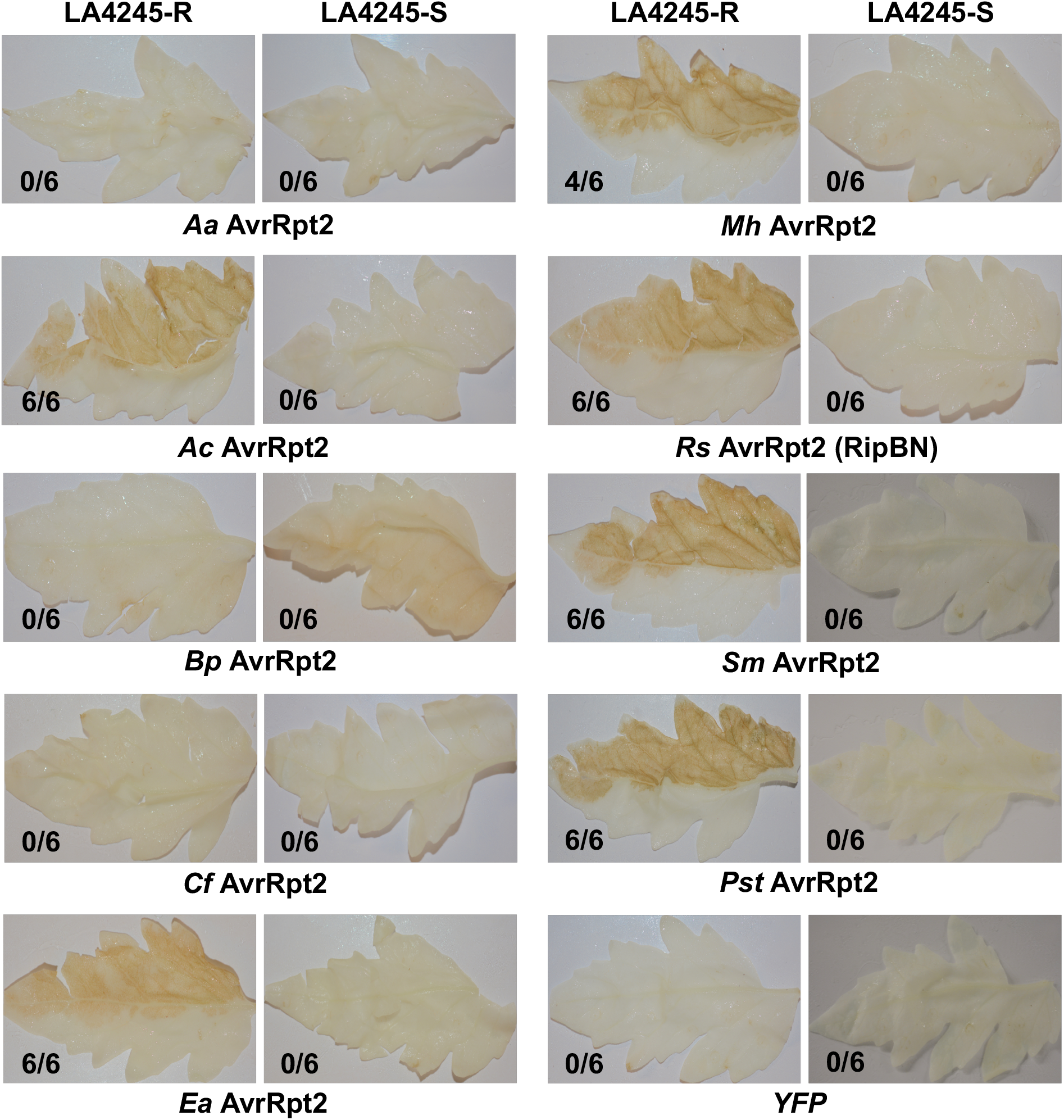
AvrRpt2 homologs from diverse bacteria are recognized by *Ptr1*. Agroinfiltration of each *avrRpt2* homolog (*Aa, A. avenae; Ac, A. citrulli; Bp, B. pyrrocinia; Cf, C. fungivorans; Ea, E. amylovora; Mh, M. huakuii; Rs, R. pseudosolanacearum; Sm, S. medicae; Pst, P. syringae* pv *tomato*) and a control (yellow fluorescent protein (YFP)) into LA4245-R and LA4245-S leaves. Detached leaves were cleared with ethanol to better visualize cell death associated with AvrRpt2 recognition. Photographs were taken four days after infiltration. Shown are the number of times cell death was observed over the total number of agroinfiltrations performed.

### An amino acid substitution in AvrRpt2 that abolishes its recognition by the apple Mr5 NLR resistance protein does not affect recognition by Ptr1

Mr5 is an NLR fire blight resistance protein in apple that recognizes strains of *Erwinia amylovora* that express AvrRpt2 (Fahrentrapp *et al*., 2012; Vogt *et al*., 2013). A single amino acid substitution in AvrRpt2 at position 156 (cysteine-to-serine) abolishes recognition of the effector by *Mr5*. In AvrRpt2NY15125 the comparable residue is tyrosine-191. Since a Y191C substitution in AvrRpt2 was shown previously to be recognized by *Ptr1* (Fig. **3**), we asked whether a Y191S substitution in AvrRpt2 would abolish *Ptr1* recognition as it does for *Mr5* recognition. Inoculation of LA4245-R plants with a DC3000 strain expressing AvrRpt2(Y191S) revealed this variant is recognized by *Ptr1*, as it induced disease resistance and reduced bacterial growth compared to an empty vector DC3000 control strain (Fig. **6a,b**). *Mr5* and *Ptr1* therefore appear to use different mechanisms to detect AvrRpt2.

**Fig. 6.**
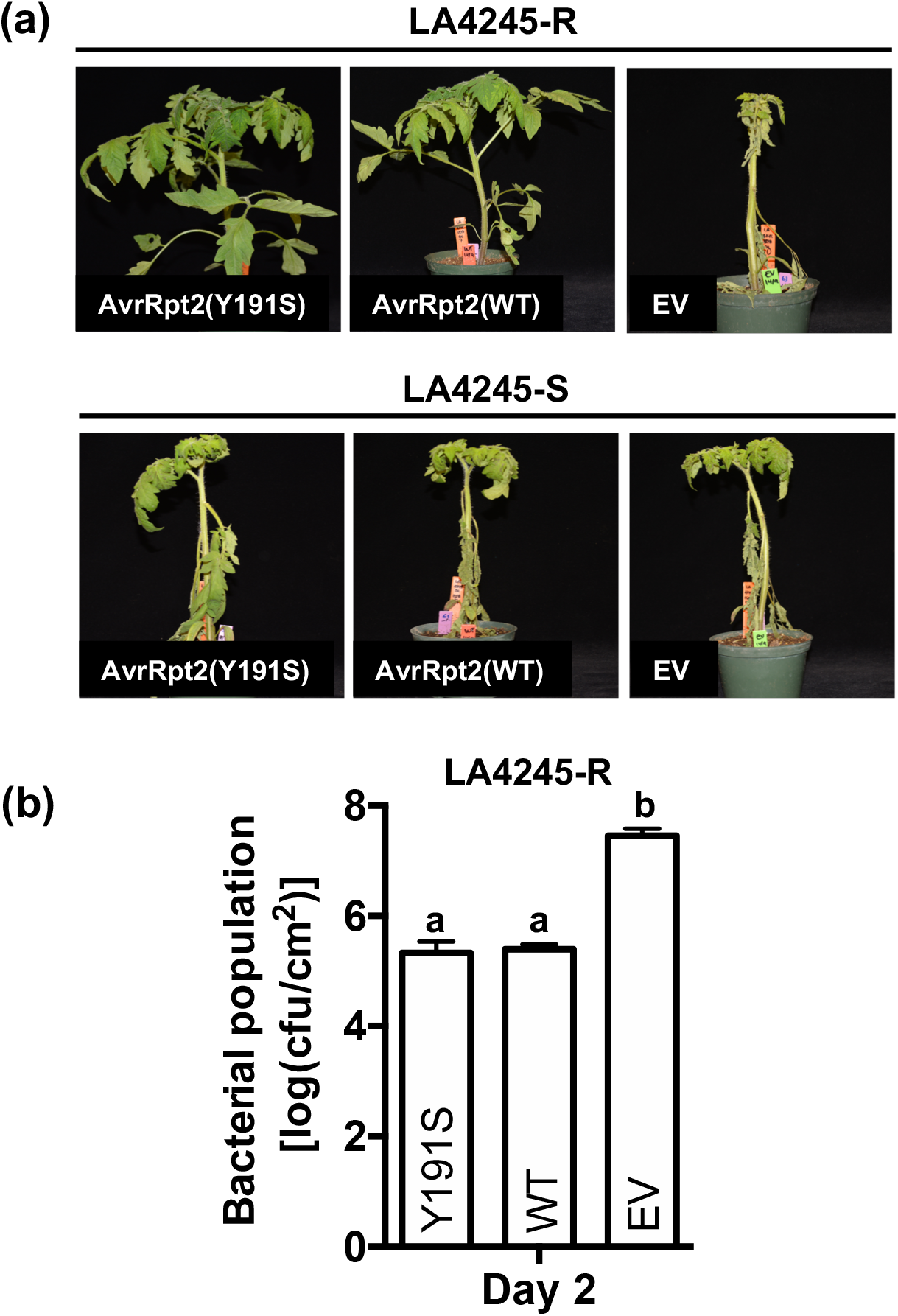
*Pst* AvrRpt2(Y191S) variant, analog of the *E. amylovora* AvrRpt2(C156S) virulent variant is recognized by *Ptr1*. (a) Disease symptoms of LA4245-R and LA4245-S plants four days after inoculation with DC3000 expressing *avrRpt2(Y191S), avrRpt2* wild-type (WT) or an empty vector (EV) at 5 x 10^4^ cfu ml^−1^. (b) Bacterial populations were measured in LA4245-R plants two days after inoculation. Significance was determined using ANOVA with a Tukey’s post hoc multiple comparison test, and different letters indicate significant differences between treatments (*P*<0.001). Bars indicate the mean of three plants and error bars represent +/− SD. Results shown are representative of three independent experiments.

### *Ptr1* confers resistance to bacterial wilt disease caused by *Ralstonia pseudosolanacearum*

The agroinfiltration experiments indicated that RipBN, the AvrRpt2 homolog from *R. pseudosolanacearum*, is recognized by *Ptr1* (Fig. **5**). The *ripBN* gene is present in a strain of *R pseudosolanacearum*, CMR15, which was collected from tomato in Cameroon in 2009 (Mahbou Somo Toukam *et al*., 2009) and later sequenced (Remenant *et al*., 2010). Bacterial wilt, caused by *R. pseudosolanacearum*, is a devastating disease for which no NLR-mediated resistance in tomato has been reported (Huet, 2014). We therefore soil drench-inoculated LA4245-R and LA4245-S plants with CMR15 and scored the number of plants showing symptoms of bacterial wilt over a 13-day time period. Beginning at 6 days after inoculation LA4245-S plants started to wilt, and 13 days after inoculation ~70% of the plants were dead (Fig. **7a,b**). Remarkably, at this timepoint there was 100% survival of the LA4245-R plants. *Ptr1*-mediated resistance therefore can be effective against bacterial wilt and might be useful for controlling bacterial wilt disease in tomato-growing areas that have *R. pseudosolanacearum* strains expressing RipBN.

**Fig. 7.**
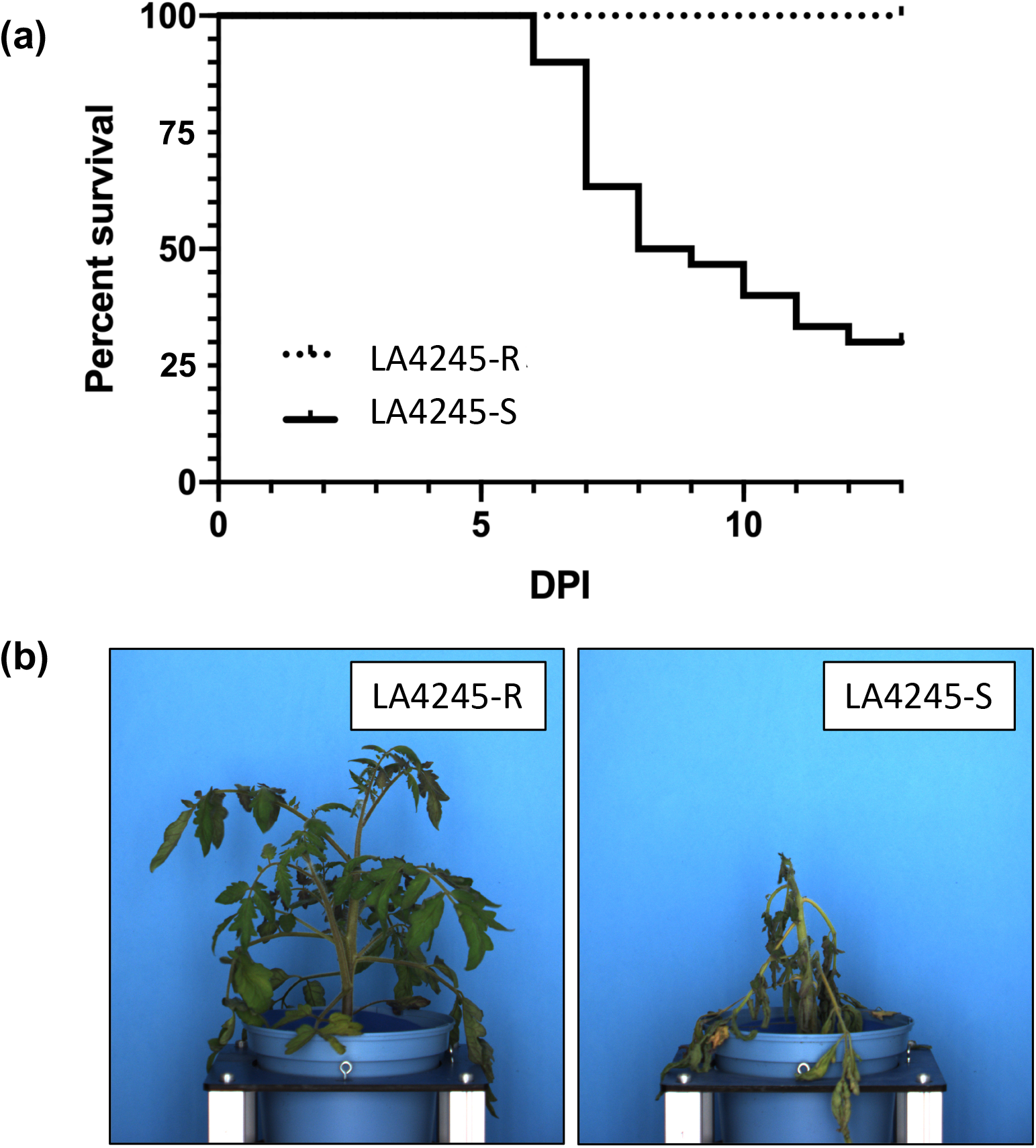
*Ptr1* confers resistance to *Ralstonia pseudosolanacearum* strain CMR15 expressing AvrRpt2 homolog RipBN. (a) LA4245-R plants (n=25) and LA4245-S plants (n=30) were soil drench-inoculated with 50 mL of 10^8^ cfu ml^−1^ of *Ralstonia pseudosolanaceraum* CMR15. The percentage of plants surviving (showing no wilt symptoms) or dying (showing severe wilting) 1 to 13 days post-inoculation (DPI) is shown. The two survival curves are significantly different at *P*<0.0001 using the Mantel-Cox test. (b) Photographs taken 13 days after inoculation show extreme wilting of a LA4245-R (left) plant and no wilting of a LA4245-S plant (right). Results shown are representative of three independent experiments.

### No RPS2, Mr5 or RIN4 orthologs are present in the *S. lycopersicoides* chromosome 4 introgression segments of LA4245-R

To initiate the map-based cloning of *Ptr1*, we generated sequence data at 14X coverage of the Heinz 1706 reference genome from genomic DNA of LA4245-R using an Illumina HiSeq2000. The reads were mapped to the *S. lycopersicum* Heinz 1706 genome sequence to identify areas of high sequence polymorphism. This analysis revealed that LA4245-R contains two *S. lycopersicoides* introgression segments on chromosome 4 in the background of the tomato parent VF36. One small segment lies within coordinates 1-260,000 bp (260 kb) and the other lies between coordinates 4,480,000 bp and 62,030,000 bp (~57.5 mb) (Fig. S8a). A high-quality genome sequence of *S. lycopersicoides* LA2951, a parent of the introgression lines, has been generated recently (see Methods). Synteny between Heinz 1706 and LA2951 was determined and gene models predicted to be in the introgressed regions of LA4245-R were identified in the *S. lycopersicoides* annotation*. Ptr1* recognition of AvrRpt2 suggests it likely encodes an NLR protein or possibly a guardee such as Rin4. No NLR-encoding genes are annotated within the small introgression segment and just 14 NLR genes are annotated within the large *S. lycopersicoides* introgression segment. Interestingly, none of these 14 genes encode a predicted protein with similarity to RPS2 or Mr5 (Fig. S8b). In fact, the *S. lycopersicoides* genes with predicted proteins having the highest similarity to RPS2 and Mr5 are all located on other chromosomes (Table S6). In addition, none of the four *Rin4* genes in tomato are located on chromosome 4 (Table S5).

## Discussion

From a serendipitous observation during a natural occurrence of bacterial speck disease we identified and further characterized two *S. lycopersicoides* introgression lines (ILs) that have strong resistance to multiple race 1 strains of *Pst*. Such resistance is important since race 1 *Pst* strains are becoming increasingly common throughout the world and yet no simply-inherited genetic resistance to these strains is known. The two ILs contain a large overlapping introgression segment from *S. lycopersicoides* chromosome 4 which carries a putative locus, which we refer to as *<u>P</u>seudomonas <u>t</u>omato <u>r</u>ace 1* (*Ptr1*), that recognizes the effector AvrRpt2. This effector is present in many *Pst* strains collected from diverse tomato-growing regions. The *Ptr1* locus, if combined with *Pto* in the same tomato variety, has the potential to become an important component of bacterial speck disease control. Here we compare the activity of *Ptr1* to Arabidopsis *RPS2* and apple *Mr5*, two genes that also detect AvrRpt2, discuss the potential utility of *Ptr1* and approaches to identifying the *Ptr1* gene, and we propose the definition of additional *Pst* races to account for the fact that two NTI loci are now known to confer resistance to bacterial speck disease.

Cleavage of Rin4 by AvrRpt2 in Arabidopsis leads to activation of RPS2 and resistance to *P. syringae* pv. tomato, although the specific mechanism by which Rin4 degradation activates RPS2 is unknown (Day *et al*., 2005; Toruño *et al*., 2018). Our analysis of AvrRpt2 variants revealed a perfect correlation between the ability of a variant to degrade tomato Rin4 proteins and its recognition by *Ptr1* and *RPS2;* AvrRpt2 variants that do not degrade Rin4 are not recognized by *Ptr1* or *RPS2*. It is possible therefore that the Ptr1 and RPS2 proteins are activated via the same mechanism subsequent to Rin4 degradation, although the possibility that a mechanism specific to tomato or Arabidopsis exists cannot currently be ruled out. The apple Mr5 protein detects activity of AvrRpt2 from *Erwinia amylovora* (Ea). The AvrRpt2_Ea_ effector does not induce Arabidopsis Rin4 degradation when the proteins are transiently co-expressed in *N. benthamiana*, however, whether Rin4 degradation in apple is correlated with Mr5 activation has not been reported (Vogt *et al*., 2013). Interestingly, an AvrRpt2_Ea_ variant that is not detected by *Mr5* is detected by *Ptr1* and is able to degrade Arabidopsis RIN4, suggesting that the recognition mechanism of these two proteins is different.

Using an anti-AtRin4 antibody, we observed that all three tomato Rin4-like proteins that are expressed in leaves were degraded in the presence of AvrRpt2. Soybean also has four Rin4 proteins and virus-induced gene silencing experiments showed that just two of them, GmRIN4a and GmRIN4b, play a role in the HR induced by AvrB and the NLR protein Rpg1b (Selote & Kachroo, 2010). A subsequent study showed that over-expression of any one of the four soybean RIN4s with Rpg1b/AvrB in leaves of *Nicotiana glutinosa* caused an HR; the authors concluded that the expression level differences in these two studies might account for these different observations (Kessens *et al*., 2014). Apple contains two Rin4-like genes although, as noted above, any specific roles they might have in activation of Mr5 have not been reported yet (Fahrentrapp *et al*., 2012; Vogt *et al*., 2013). In the future, we will investigate the requirement of the three tomato Rin4 proteins for activation of Ptr1 using both CRISPR-generated mutations in the corresponding genes and, once the *Ptr1* gene is identified, transient co-expression in *N. benthamiana*.

We found that *Ptr1* is able to recognize AvrRpt2 homologs from a diverse array of different bacteria including several plant pathogens. Each of the AvrRpt2 proteins recognized by *Ptr1* had been previously shown to also to induce *At*Rin4 degradation in Arabidopsis, although their ability to activate RPS2-mediated resistance was not reported (Eschen-Lippold *et al*., 2016). If *avrRpt2* homologs are widespread in field isolates of *Acidovorax citrulli* and *Erwinia amylovora* then our results suggest that *Ptr1* might be useful if expressed transgenically in cucurbits to control bacterial fruit blotch caused by *A. citrulli* and in apple or pear to confer resistance to fire blight caused by *E. amylovora*. Despite extensive screening of tomato germplasm, no single *R* gene has been identified which confers resistance to bacterial wilt disease caused by *R. pseudosolanacearum* (Huet, 2014). We found that *Ptr1* was remarkably effective in preventing symptoms of bacterial wilt in a growth chamber assay using strain CMR15. Unfortunately, CMR15 seems to be a rare example of a *R. pseudosolanacearum* strain that expresses an AvrRpt2 homolog (RipBN) (Peeters *et al*., 2013) (https://iant.toulouse.inra.fr//bacteria/annotation/site/prj/T3Ev3/), and *Ptr1* will therefore likely not be of broad utility for controlling bacterial wilt disease. Nevertheless, further study of interaction between CMR15 and LA4245-R might provide some useful insights into the molecular basis of the NTI response against *R. pseudosolanacearum* that is induced by *Ptr1*.

Considering that *Ptr1* plays a role in detecting the type III effector AvrRpt2 it seems most likely that it encodes an NLR, although it could encode a decoy or guardee protein which is monitored by an NLR. In either case, one of these genes must be located in the introgression segments although the other one may or may not lie in the introgressions. Examples of each possibility are known: the *Pto* (decoy) and *Prf* (NLR) genes are located in a 20 kb region of chromosome 5, whereas the *RIN4* (guardee) and *RPS2* (NLR) genes are located on different chromosome in Arabidopsis. In the Heinz 1706 tomato reference genome sequence chromosome 4 has 56 NLR-encoding genes, the most of any chromosome; 43 of these genes are clustered tightly within a 100-kilobase region with the other 13 located throughout the rest of the chromosome. *S. lycopersicoides* has 66 NLR-encoding genes on chromosome 4 with a similar distribution as seen in Heinz 1706 – 58 are tightly clustered at one end and 8 are distributed along the chromosome.

One boundary of the large introgression segment occurs in the middle of the large NLR cluster which potentially eliminates as *Ptr1* candidates 51 NLR-encoding genes. Each of the remaining 15 NLR genes is a candidate for *Ptr1*. None of these candidates encode proteins with obvious similarity to RPS2 or Mr5, and if one of these is Ptr1, it will be just the third known example of convergent evolution in different plant species for recognition of the same effector (Ashfield *et al*., 2004; Carter *et al*., 2018). The obvious decoy/guardee proteins that might be involved with AvrRpt2 recognition in tomato are the three Rin4-like proteins expressed in leaves. None of these genes are located on chromosome 4 of tomato or *S. lycopersicoides*. Another, perhaps less likely, scenario that should be considered is that a gene in the introgression segment encodes a novel host protein that acts with Rin4 to activate an NLR located on another chromosome in VF36. All of these possibilities will be investigated in the future as we seek to identify the genes involved in AvrRpt2 recognition and understand the associated molecular mechanisms.

Although LA4245-R plants are morphologically very similar to VF36, the IL line has two introgression segments on chromosome 4, including one large (~57.5 mb) segment encompassing the majority of the 66.56 mb chromosome; however, this IL line cannot be maintained in a homozygous condition. If *Ptr1* is to be useful for control of speck disease, it will need to be introgressed into other tomato breeding lines and it will be necessary to reduce the size of the introgressed segment in order to reduce ‘linkage drag’ (i.e., deleterious alleles). However, recombination is often severely suppressed in plants carrying chromosomal regions from a wild relative of tomato, and *S. lycopersicoides* is particularly distant from cultivated tomato (Grandillo *et al*., 2011). One approach that might be used in this case is to cross LA4245-R with a species which is phylogenetically intermediate between tomato and *S. lycopersicoides*. A good candidate for such a ‘bridge’ species is *S. pennellii* which has been shown previously to be useful in increasing recombination frequency in *S. lycopersicoides* introgression regions (Canady *et al*., 2006). Importantly, *Ptr1* and *Pto* are located on different chromosomes (4 and 5, respectively) which will facilitate introgressing them both into advanced breeding lines.

The identification of a second *R* gene that confers resistance to bacterial speck disease presents the opportunity to extend the currently defined race structure of *P. syringae* pv. tomato. The two current races were defined based on the *Pto* gene in which race 0 strains express either or both of the type III effectors AvrPto and AvrPtoB and race 1 strains evade *Pto* detection by either losing or mutating these effector genes, or in the case of AvrPtoB suppressing its protein accumulation (Pedley & Martin, 2003; Lin *et al*., 2006; Kunkeaw *et al*., 2010). Based on the discovery of *Ptr1*, and focusing on *Pst* strains related to T1, we propose that race 0 strains might be considered the original state of the pathogen and refer to those strains that express *avrPto* or *avrPtoB* along with *avrRpt2* (Table 1). Race 1 can then refer to strains that express *avrRpt2*, but lack *avrPto* or *avrPtoB*. A newly defined race 2 would refer to strains that have *avrPto* or *avrPtoB*, but lack *avrRpt2*. Finally, a hypothetical strain that lacks all three of these effectors would be defined as race 3. Examples of race 0, 1, and 2 *Pst* strains are provided in Table 1 along with the ability of *Ptr1* and/or *Pto* to recognize them.

**Table 1.**
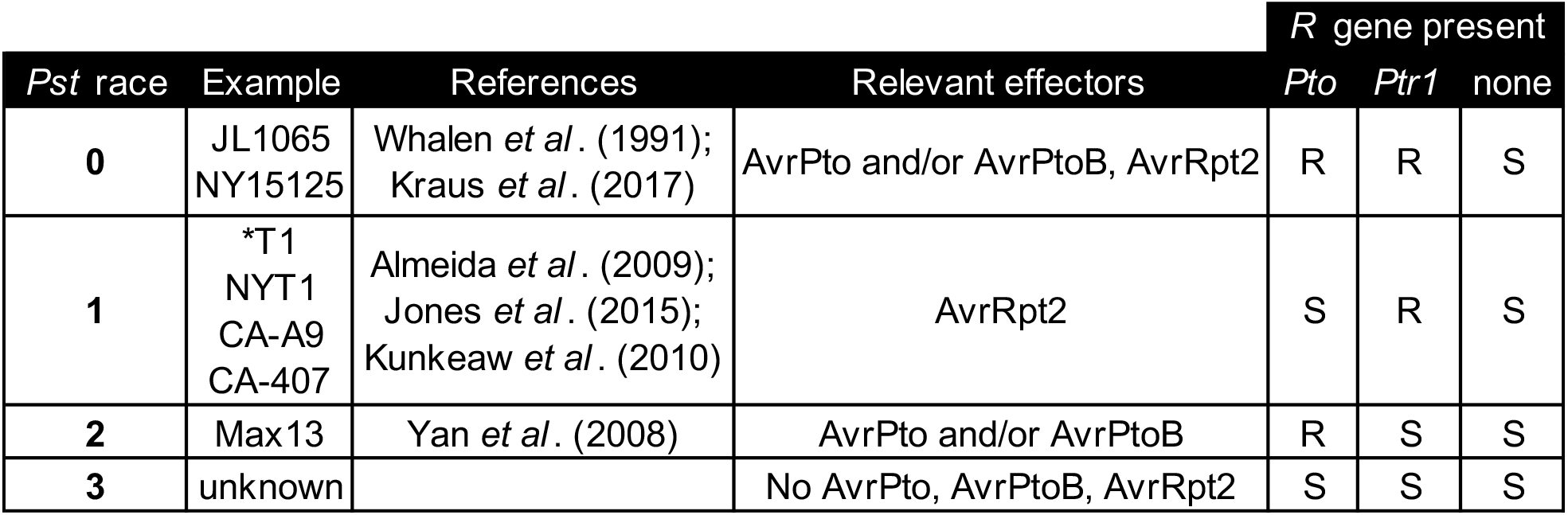
Proposed races for *P. syringae* pv. tomato based on the *Pto* and *Ptr1* genes. Resistance and susceptibility are denoted as R and S, respectively. *Race 1 strains T1, NYT1, CA-A9 and CA-407 are positive for the presence of the *avrPtoB* gene, but protein does not accumulate for unknown reasons.

|  |  |  |  | R gene present |  |  |
| --- | --- | --- | --- | --- | --- | --- |
| *Pst* race | Example | References | Relevant effectors | *Pto* | *Ptr1* | none |
| **0** | JL1065<br>NY15125 | Whalen *et al.* (1991);<br>Kraus *et al.* (2017) | AvrPto and/or AvrPtoB, AvrRpt2 | R | R | S |
| **1** | *T1<br>NYT1<br>CA-A9<br>CA-407 | Almeida *et al.* (2009);<br>Jones *et al.* (2015);<br>Kunkeaw *et al.* (2010) | AvrRpt2 | S | R | S |
| **2** | Max13 | Yan *et al.* (2008) | AvrPto and/or AvrPtoB | R | S | S |
| **3** | unknown |  | No AvrPto, AvrPtoB, AvrRpt2 | S | S | S |

In summary, *Ptr1* has the potential to become an important component (along with *Pto*) for the control of bacterial speck disease in tomato. In addition, the eventual cloning and characterization of the gene will allow its molecular characterization and, in particular, might shed light on whether it uses the same or a different mechanism than used by RPS2 and Mr5 to respond to AvrRpt2-mediated degradation of Rin4.

## Acknowledgments

We thank Robyn Roberts and Alan Collmer for helpful comments on the manuscript, Justin Lee for the AvrRpt2 homologs, Barbara Kunkel for JL1065 strains, Gitta Coaker for anti-RIN4 antibodies, Dan Klessig for *A. thaliana rps2* seeds, Kathy Munkvold and Fanhong Meng for generating DC3000 strains with T1 effectors, and Roger Chetelat and the Tomato Genetics Resource Center for seeds of LA4245. The *Ralstonia* assays were performed at the Toulouse Plant-Microbe Phenotyping facility of the LIPM – UMR (INRA441/CNRS2594). This work was supported by National Science Foundation grant IOS-1546625 (GBM), NSF Research Experience for Undergraduates grant DBI-1358843 (MV), Laboratoire d’Excellence (LABEX), TULIP (ANR-10-LABX-41; NP, FMM), NSF grant IOS-0923312 (JJG), and Colciencias grant 673 (CMM).

## Author contributions

Conceived and designed the experiments CMM, SRH, GBM. Performed the experiments: CMM, SM, SRH, MV, CMK, FMM and NP. Analyzed the data: CMM, GBM. Contributed reagents/materials/data analysis: ML, SS, SRS, AF, JGG, CDS, and NP. Wrote the paper: CMM and GBM. All authors read and approved the manuscript.

## Supporting Information

**Fig. S1** Comparison of the *Pst* NY15125 chromosome with the chromosomes of *Pst* strains DC3000 and T1.

**Fig. S2** Deletion of *avrRpt2* from *Pst* strain JL1065 abolishes recognition by *Ptr1*.

**Fig. S3** Ptr1 confers resistance to several race 1 *Pseudomonas syringae* pv. tomato strains.

**Fig. S4** Immunoblotting confirms expression of each AvrRpt2 variant in *Pst* DC3000.

**Fig. S5** DC3000 strains expressing AvrRpt2 or the AvrRpt2 variants cause similar disease symptoms and grow to the same levels in LA4245-S plants.

**Fig. S6** AvrRpt2 variants F70R, E150S, Y191C and D216E are recognized by *RPS2*.

**Fig. S7** *Agrobacterium-mediated* transient expression of AvrRpt2 homologs in *Nicotiana benthamiana* leaves.

**Fig. S8** The predicted proteins of the 15 NLR-encoding genes in the introgressed segment of LA4245 bear little similarity to RPS2 or Mr5.

**Table S1** Bacterial strains used in this study.

**Table S2** Vectors and plasmids used in this study.

**Table S3** Oligonucleotides used in this study.

**Table S4** Summary of the type III effectors present in the P. *syringae* strains NY15125, DC3000, T1, NYT1, CA-407, and CA-A9.

**Table S5** Features and expression patterns of the four genes in tomato that encode proteins with similarity to Arabidopsis RIN4.

**Table S6** Gene models with the highest similarity to RPS2 and Mr5 in *S. lycopersicoides*.

