## Supplementary material for "The *Ptr1* locus of *Solanum lycopersicoides* confers resistance to race 1 strains of *Pseudomonas syringae* pv. tomato and to *Ralstonia pseudosolanacearum* by recognizing the type III effectors AvrRpt2/RipBN": Suplemental Information

**Supplemental Figures**

**Fig. S1** Comparison of the *Pst* NY15125 chromosome with the chromosomes of *Pst* strains DC3000 and T1.

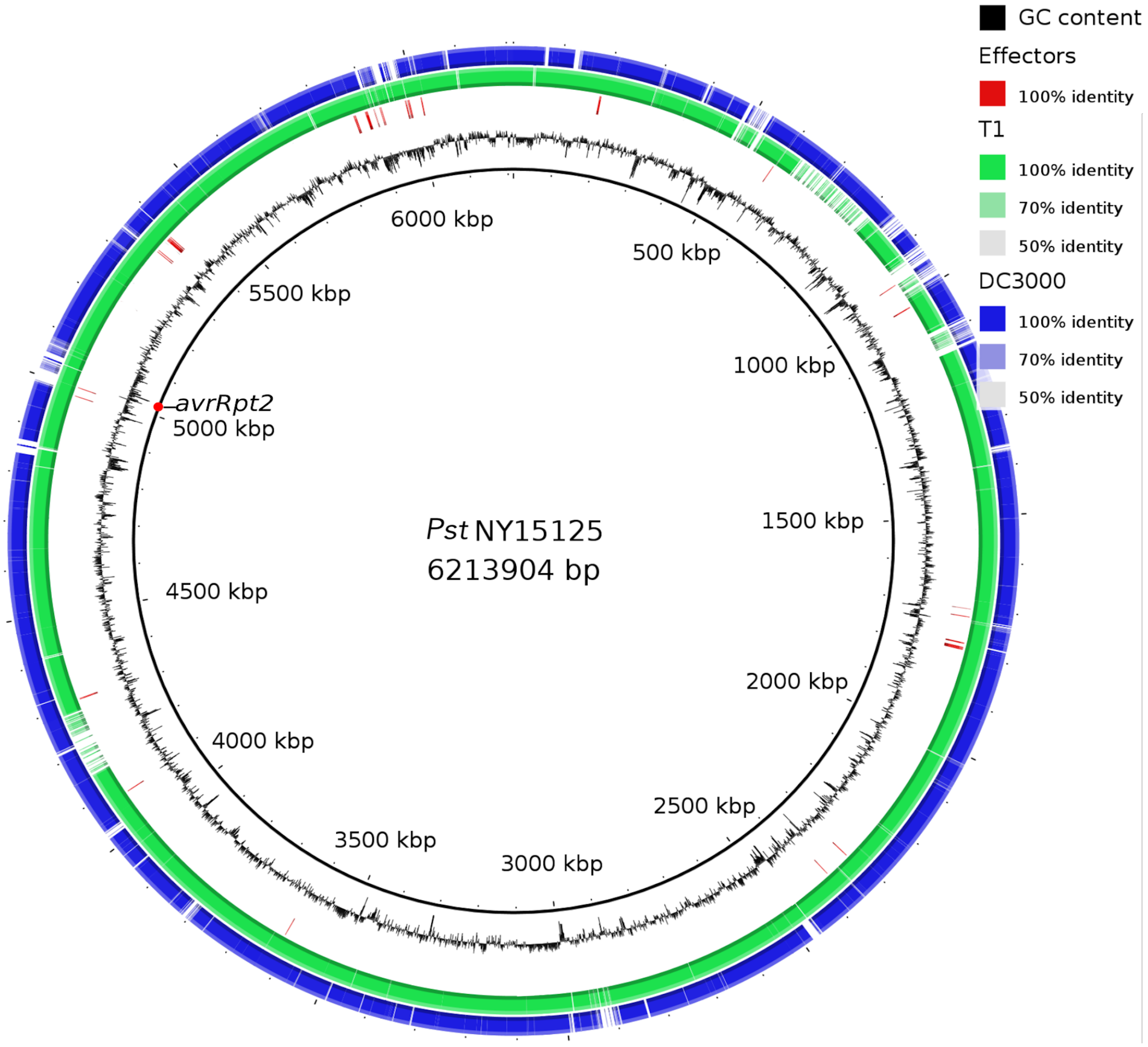


The solid black line represents the NY15125 chromosome and the jagged black line represents GC percentage. The locations of the type III effectors are shown with short red lines, and location of *avrRpt2* is shown with a red dot. Genome comparisons between NY15125 and T1 or DC3000 are depicted in blue or green, respectively. Ring image was generated using the BLAST Ring Image Generator (BRIG) http://sourceforge.net/projects/brig/.

**Fig. S2** Deletion of *avrRpt2* from *Pst* strain JL1065 abolishes recognition by *Ptr1*.

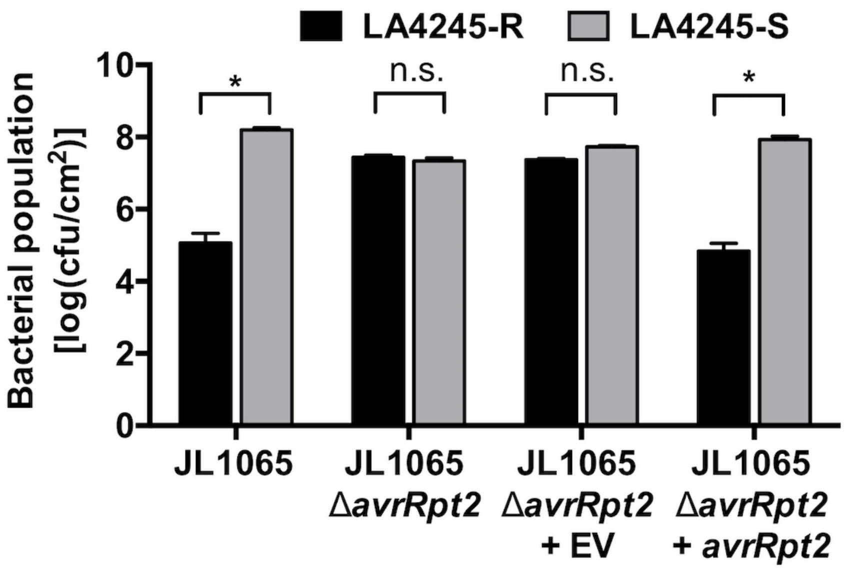


LA4245-R and LA4245-S plants inoculated with JL1065 wild-type, JL1065Δ*avrRpt2,* a complemented strain, or an empty vector (EV) strain at 1 x $10^4$ cfu $ml^{-1}$. Bacterial populations were measured four days after inoculation. Significance was determined by a pair-wise t-test and is indicated as: * $P$<0.05 or not significant (n.s.) at $P$>0.05. Bars indicate the mean of three independent experiments using three plants per strain. Error bars represent +/- SEM.

**Fig. S3** Ptr1 confers resistance to several race 1 *Pseudomonas syringae* pv. tomato strains.

| ***P. syringae* pv. tomato CA-A9** | ***P. syringae* pv. tomato CA-407** | ***P. syringae* pv. tomato** |
|---|---|---|

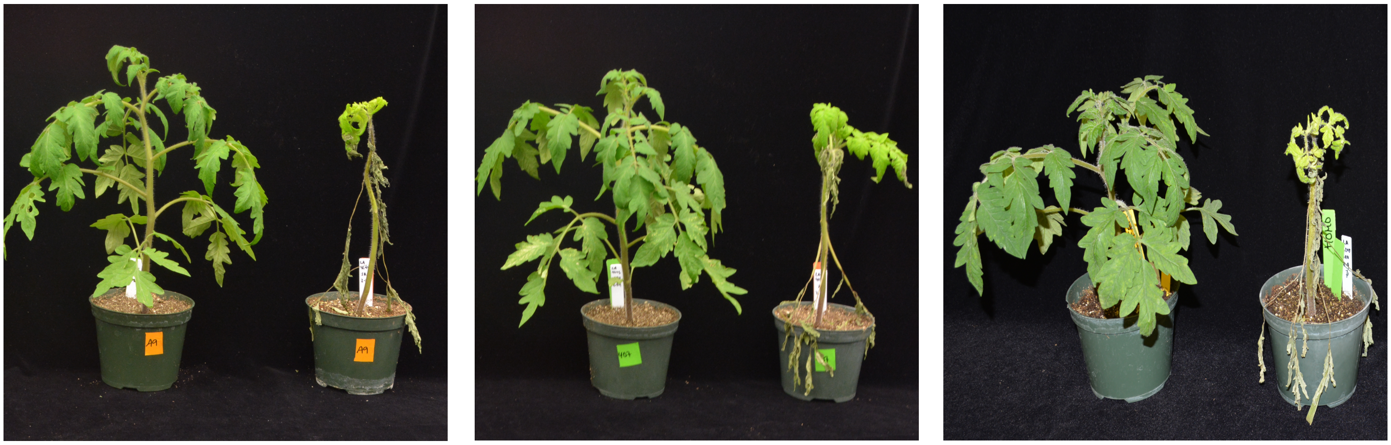

| LA4245-R | LA4245-S | LA4245-R | LA4245-S | LA4245-R | LA4245-S |
|---|---|---|---|---|---|

Disease symptoms of LA4245-R and LA4245-S plants vacuum infiltrated with *Pst* strains CA-A9, CA-407 and NYT1 at 1 x $10^4$ cfu $ml^{-1}$ and photographed eight days later.

**Fig. S4** Immunoblotting confirms expression of each AvrRpt2 variant in *Pst* DC3000.

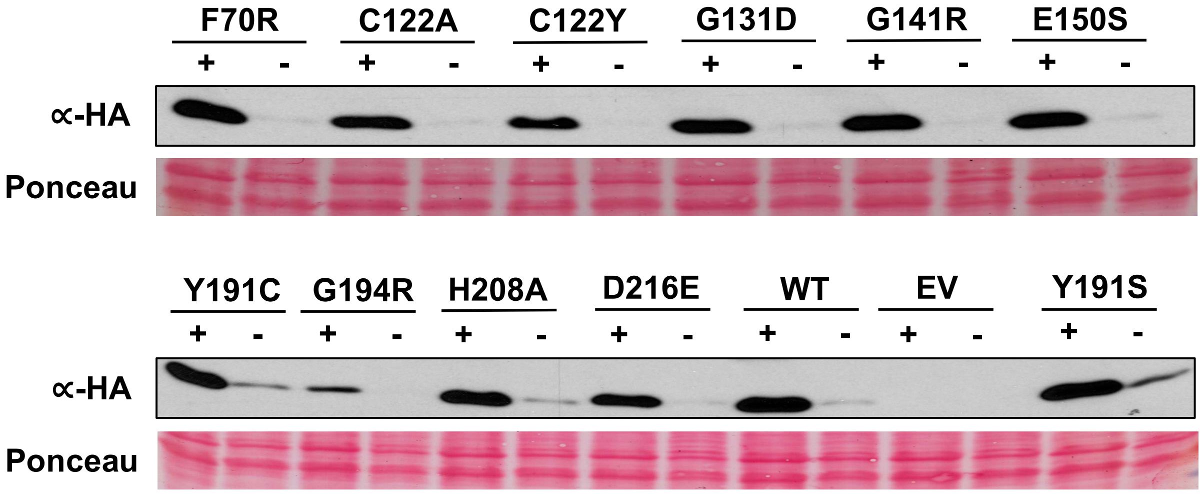


Detection of protein accumulation of each AvrRpt2 variant by immunoblotting using ∝-HA. Bacteria were grown in *hrp*-inducing minimal media (+) or KB media (-) to detect each effector protein. As expected, proteins were detectable only when the strains were grown in *hrp*-inducing medium (+). Ponceau staining shows amount of protein loaded in each lane.

**Fig. S5** DC3000 strains expressing AvrRpt2 or the AvrRpt2 variants cause similar disease symptoms and grow to the same levels in LA4245-S plants.

(a)

(b)

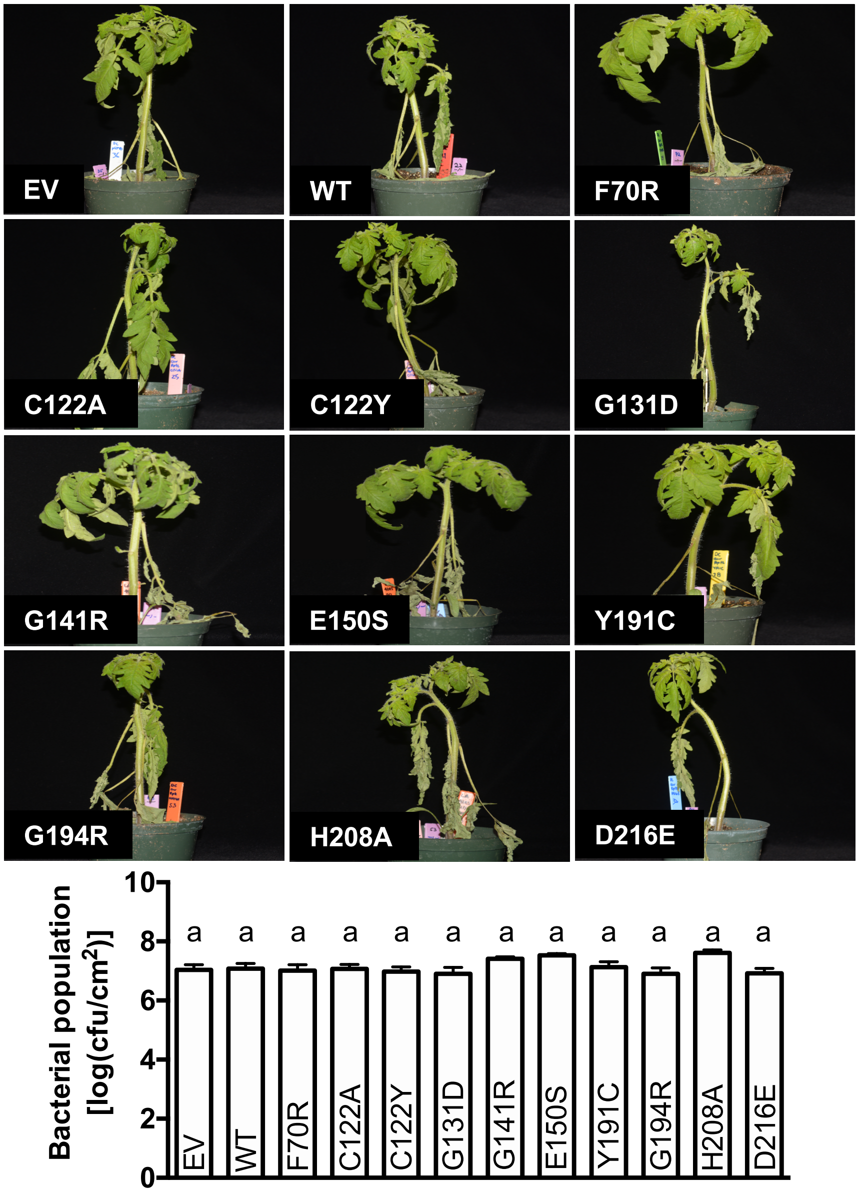


(a) Whole-plant symptoms of LA4245-S plants inoculated with DC3000 carrying *avrRpt2* wild-type (WT), *avrRpt2* variants or an empty vector (EV) at 5 x $10^4$ cfu $ml^{-1}$ five days after inoculation. (b) Bacterial populations in leaves of LA4245-S plants inoculated with DC3000 expressing *avrRpt2* wild-type (WT), *avrRpt2* variants or an empty vector (EV), 2 days after inoculation. Significance was determined using ANOVA with a Tukey´s post hoc multiple comparison test, and different letters indicate significant differences between treatments ($P$<0.001). Bars indicate the mean of three independent experiments using three plants per strain. Error bars represent +/- SEM

**Fig. S6** AvrRpt2 variants F70R, E150S, Y191C and D216E are recognized by *RPS2*.

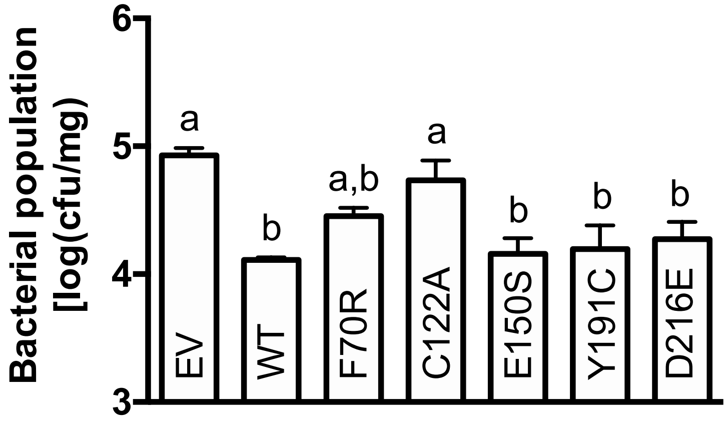


Five week-old *Arabidopsis* Col-0 *RPS2* plants were dip inoculated with DC3000 carrying *avrRpt2* wild-type (WT), *avrRpt2* variants or an empty vector (EV) at 3 x $10^8$ cfu $ml^{-1}$. Bacterial populations were measured three days after inoculation. Significance was determined using ANOVA with a Tukey´s post hoc multiple comparison test, and different letters indicate significant differences between treatments ($P$<0.1). Bars indicate the mean of three plants and error bars represent +/- SEM. Data are representative of three independent experiments.

**Fig. S7** *Agrobacterium*-mediated transient expression of AvrRpt2 homologs in *Nicotiana benthamiana* leaves.

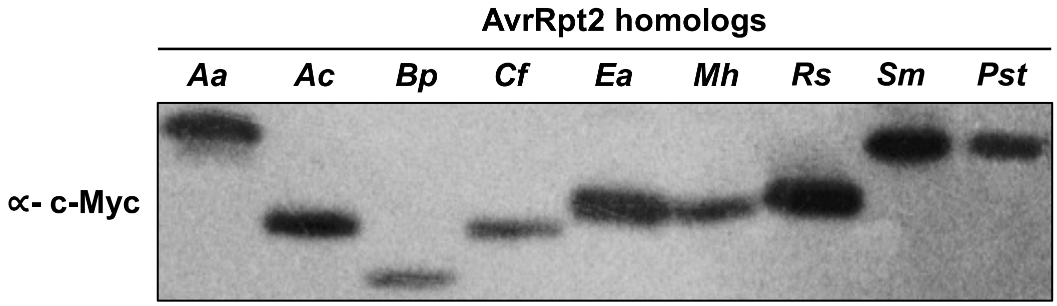


Agroinfiltration of AvrRpt2 homologs (*Aa, A. avenae; Ac, A. citrulli; Bp, B. pyrrocinia; Cf, C. fungivorans; Ea, E. amylovora; Mh, M. huakuii; Rs, R. pseudosolanacearum; Sm, S. medicae; Pst, P. syringae* pv *tomato)* at an $OD_{600}$ of 0.3 into *N. benthamiana* leaves. Samples were taken 43 hours after infiltration. Total proteins extracted from infiltrated leaves were subjected to immunoblotting using an ∝-c-Myc antibody.

**Fig. S8** The predicted proteins of the 15 NLR-encoding genes in the introgressed segment of LA4245 bear little similarity to RPS2 or Mr5.

(a)

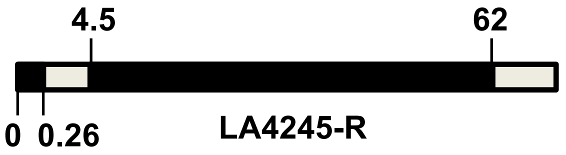


(b)

| Gene model | % similarity to RPS2 | % similarity to Mr5 | No. amino acid | Annotation | | |
|---|---|---|---|---|---|---|
| Solyd04g057440 | 47.8 | 34.7 | 911 | CC | NB-ARC | LRR |
| Solyd04g057510 | 47.3 | 33.9 | 870 | | NB-ARC | LRR |
| Solyd04g057520 | 47.8 | 34.5 | 908 | CC | NB-ARC | LRR |
| Solyd04g057570 | 52.8 | 56.7 | 452 | | NB-ARC | |
| Solyd04g057630 | 49.3 | 44.8 | 718 | | NB-ARC | LRR |
| Solyd04g057640 | 49.9 | 56.4 | 391 | CC | NB-ARC | |
| Solyd04g058020 | 43.6 | 39.9 | 655 | | NB-ARC | LRR |
| Solyd04g059470 | 48.8 | 39.4 | 733 | | NB-ARC | LRR |
| Solyd04g059610 | 53.8 | 56.4 | 850 | CC | NB-ARC | LRR |
| Solyd04g060430 | 40.6 | 39.3 | 424 | CC | NB-ARC | |
| Solyd04g060640 | 49.7 | 45.8 | 795 | CC | NB-ARC | LRR |
| Solyd04g061490 | 54.2 | 42.9 | 593 | | NB-ARC | LRR |
| Solyd04g064750 | 43.4 | 32.4 | 1073 | CC | NB-ARC | LRR |
| Solyd04g067320 | 40.3 | 36.8 | 1159 | TIR | NB-ARC | LRR |
| Solyd04g076500 | 42.7 | 30.7 | 820 | | NB-ARC | LRR |

(a) Scheme of the introgressed segment present in LA4245-R. Coordinates (in megabases, Mb) are based on SNP density mapped to *S. lycopersicum* Heinz 1706 SL2.50. (b) Predicted NLR-like genes present in the large introgressed segment of LA4245-R (no NLR genes are present in the small introgressed segment). Gray indicates absence of this domain.

**Supplemental Tables**

**Table S1** Bacterial strains used in this study.

| Name | Genotype | Race | Source | Relevant efectors |
|---|---|---|---|---|
| *P. syringae* pv. tomato DC3000 | Wild type | 0 | (Buell *et al.*, 2003) | AvrPto |
| *P. syringae* pv. tomato NY15125 | Wild type | 0 | (Kraus *et al.*, 2017) | AvrPto, AvrRpt2 |
| *P. syringae* pv. tomato JL1065 | Wild type | 0 | (Whalen *et al.*, 1991) | AvrPto, AvrRpt2 |
| *P. syringae* pv. tomato T1 | Wild type | 1 | (Almeida *et al.*, 2009) | AvrRpt2 |
| *P. syringae* pv. tomato NYT1 | Wild type | 1 | (Jones *et al.*, 2015) | AvrRpt2 |
| *P. syringae* pv. tomato CA-A9 | Wild type | 1 | (Kunkeaw *et al.*, 2010) | AvrRpt2 |
| *P. syringae* pv. tomato CA-407 | Wild type | 1 | (Kunkeaw *et al.*, 2010) | AvrRpt2 |
| *R. pseudosolanacearum* CMR15 | Wild type | - | (Mahbou Somo Toukam *et al.*, 2009) | RipBN |
| *A. tumefaciens* 1D1249 | Wild type | - | (Wroblewski *et al.*, 2005) | - |
| *E.coli* TOP 10 | Wild type | - | Thermo Fisher Co. | - |
| *E.coli* S17-1 | Wild type | - | A. Collmer, Cornell University | - |

**Table S2** Vectors and plasmids used in this study.

| Vector | Insert | Identifier | Purpose | Source |
|---|---|---|---|---|
| pK18mobsac | - | - | Biparental Mating | (Kvitko & Collmer, 2011) |
| pK18mobsac | Δ*avrRpt2* | pCM3 | For deletion of *avrRpt2* in NY15125 by biparental mating | This work |
| pENTR SD TOPO | - | - | Gateway entry vector | Thermo Fisher |
| pENTR SD TOPO | *avrRpt2* (no stop) | 7208 | For LR recombination into destination vector | This work |
| pENTR SD TOPO | *avrRpt2*F70R (no stop) | pCM4 | For LR recombination into destination vector | This work |
| pENTR SD TOPO | *avrRpt2*C122A (no stop) | pCM5 | For LR recombination into destination vector | This work |
| pENTR SD TOPO | *avrRpt2*C122Y (no stop) | pCM17 | For LR recombination into destination vector | This work |
| pENTR SD TOPO | *avrRpt2*G131D (no stop) | pCM18 | For LR recombination into destination vector | This work |
| pENTR SD TOPO | *avrRpt2*G141R (no stop) | pCM30 | For LR recombination into destination vector | This work |
| pENTR SD TOPO | *avrRpt2*E150S (no stop) | pCM27 | For LR recombination into destination vector | This work |
| pENTR SD TOPO | *avrRpt2*Y191C (no stop) | pCM6 | For LR recombination into destination vector | This work |
| pENTR SD TOPO | *avrRpt2*Y191S (no stop) | pCM53 | For LR recombination into destination vector | This work |
| pENTR SD TOPO | *avrRpt2*G194R (no stop) | pCM19 | For LR recombination into destination vector | This work |
| pENTR SD TOPO | *avrRpt2*H208A (no stop) | pCM31 | For LR recombination into destination vector | This work |
| pENTR SD TOPO | *avrRpt2* D216E (no stop) | pCM7 | For LR recombination into destination vector | This work |
| pCPP5372 | - | - | Binary Gateway destination vector for type three effector protein expression with C-terminal HA-tag | (Oh *et al.*, 2007) |
| pCPP5372 | *avrRpt2* | 7200 | Generating hrp protein fusion with C-terminal HA-tag | This work |
| pCPP5372 | *avrRpt2*F70R | pCM9 | Generating hrp protein fusion with C-terminal HA-tag | This work |
| pCPP5372 | *avrRpt2*C122A | pCM10 | Generating hrp protein fusion with C-terminal HA-tag | This work |
| pCPP5372 | *avrRpt2*C122Y | pCM20 | Generating hrp protein fusion with C-terminal HA-tag | This work |

| | | | | |
|---|---|---|---|---|
| pCPP5372 | *avrRpt2*G131D | pCM21 | Generating hrp protein fusion with C-terminal HA-tag | This work |
| pCPP5372 | *avrRpt2*G141R | pCM32 | Generating hrp protein fusion with C-terminal HA-tag | This work |
| pCPP5372 | *avrRpt2*E150S | pCM28 | Generating hrp protein fusion with C-terminal HA-tag | This work |
| pCPP5372 | *avrRpt2*Y191C | pCM11 | Generating hrp protein fusion with C-terminal HA-tag | This work |
| pCPP5373 | *avrRpt2*Y191S | pCM55 | Generating hrp protein fusion with C-terminal HA-tag | This work |
| pCPP5372 | *avrRpt2*G194R | pCM22 | Generating hrp protein fusion with C-terminal HA-tag | This work |
| pCPP5372 | *avrRpt2*H208A | pCM33 | Generating hrp protein fusion with C-terminal HA-tag | This work |
| pCPP5372 | *avrRpt2*D216E | pCM33 | Generating hrp protein fusion with C-terminal HA-tag | This work |
| pGWB417 | - | - | Binary Gateway destination vector for plant expression with C-terminal Myc-tag. | (Nakagawa *et al.*, 2007) |
| pGWB417 | *avrRpt2_Acidovorax avenae subsp. avenae* ATCC 19860 | pTK187 | Generating C-terminal Myc-tag fusion | This work |
| pGWB417 | *avrRpt2_Acidovorax citrulli* tw6 | pTK188 | Generating C-terminal Myc-tag fusion | This work |
| pGWB417 | *avrRpt2_Burkholderia pyrrocinia* Lyc2 | pTK189 | Generating C-terminal Myc-tag fusion | This work |
| pGWB417 | *avrRpt2_Collimonas fungivorans* | pTK190 | Generating C-terminal Myc-tag fusion | This work |
| pGWB417 | *avrRpt2_Erwinia amylovora* ATCC 49946 | pTK191 | Generating C-terminal Myc-tag fusion | This work |
| pGWB417 | *avrRpt2_Mesorhizobium huakuii* 7653R | pTK192 | Generating C-terminal Myc-tag fusion | This work |
| pGWB417 | *avrRpt2_Pseudomonas syringae pv. tomato* NY15125 | pCM1 | Generating C-terminal Myc-tag fusion | This work |
| pGWB417 | *avrRpt2_Ralstonia solanacearum* CMR15 | pTK194 | Generating C-terminal Myc-tag fusion | This work |
| pGWB417 | *avrRpt2_Sinorhizobium medicae* WSM1369 | pTK195 | Generating C-terminal Myc-tag fusion | This work |

**Table S3** Oligonucleotides used in this study.

| Gene | Direction | Identifier | Sequence (5'→3') | Source |
|---|---|---|---|---|
| Introgressed region in Chr4 | Forward | Spenn-ch04_5416619 | taatgaggcagagcaagttt | This work |
| Introgressed region in Chr4 | Reverse | Spenn-ch04_5416620 | ccctcaagaaccatgaatca | This work |
| *avrRpt2* | Forward | oCM26 | CACCatgaaaattgctccagttgccataaatcacag | This work |
| *avrRpt2* | Reverse | oCM27 | cacacgcaatgctctaccgc | This work |
| 5' UTR *avrRpt2* | Forward | oCM20 | atcggGAATTCgtgctgatggatgctgcagg | This work |
| 5' UTR *avrRpt2* | Reverse | oCM21 | ccacgtgaagataccttgctgcttgttaagtcgtccgttgg | This work |
| 3' UTR *avrRpt2* | Forward | oCM22 | agcaaggtatcttcacgtggcgg | This work |
| 3' UTR *avrRpt2* | Reverse | oCM23 | atcggCCCGGGttctgcgagcgatttgcggg | This work |
| UTR *avrRpt2* | Forward | oCM38 | ctgatcatgtgtgccttgacccc | This work |
| UTR *avrRpt2* | Reverse | oCM39 | ggactgcagggtgtttatcggg | This work |
| UTR *avrRpt2* | Forward | 3719 | ttagctcactcattaggcacc | This work |
| UTR *avrRpt2* | Reverse | 3720 | cctcttcgctattacgcca | This work |
| *avrRpt2* F70R | Forward | oCM28 | atagaggttccagccCGTggagggtggttc | This work |
| *avrRpt2* F70R | Reverse | oCM29 | ccctccACGggctggaacctctatcttg | This work |
| *avrRpt2* C122A | Forward | oCM30 | gcgaatgggaGCTtggtatgcctgcg | This work |
| *avrRpt2* C122A | Reverse | oCM31 | cataccaAGCtcccattcgctcattac | This work |
| *avrRpt2* C122Y | Forward | oCM74 | gcgaatgggaTATtggtatgcctgc | This work |
| *avrRpt2* C122Y | Reverse | oCM75 | gcataccaATAtcccattcgctcattacc | This work |
| *avrRpt2* G131D | Forward | oCM76 | agaatggttGACcattctgtcgaagc | This work |
| *avrRpt2* G131D | Reverse | oCM77 | agcttcgacagaatgGTCaaccattc | This work |
| *avrRpt2* G141R | Forward | oCM104 | cctaAGActgccggagctctatgagggaag | This work |
| *avrRpt2* G141R | Reverse | oCM105 | agagctccggcagTCTtaggcgaggcccagcttc | This work |
| *avrRpt2* E150S | Forward | oCM86 | ctatgagggaaggTCAggcccagctgggctac | This work |
| *avrRpt2* E150S | Reverse | oCM87 | gtagcccagctgggccTGAccttccctcatag | This work |
| *avrRpt2* Y191C | Forward | oCM32 | gtgcactgttgTGTaagcacgggccg | This work |
| *avrRpt2* Y191C | Reverse | oCM33 | ccgtgcttACAcaacagtgcacccaactc | This work |
| *avrRpt2* Y191C | Forward | oCM109 | gttgggtgcactgttgTCTaagcacg | This work |
| *avrRpt2* Y191C | Reverse | oCM110 | ccgtgcttAGAcaacagtgcaccc | This work |
| *avrRpt2* G194R | Forward | oCM80 | gttgtataagcacAGGccgattatatttggg | This work |
| *avrRpt2* G194R | Reverse | oCM81 | cccaaatataatcggCCTgtgcttatacaac | This work |
| *avrRpt2* H208A | Forward | oCM106 | gctggGCTatgtcggtcctcactggtgtcg | This work |
| *avrRpt2* H208A | Reverse | oCM107 | gaggaccgacatAGCccagctgtcattcggagttttc | This work |
| *avrRpt2* D216E | Forward | oCM34 | ctcactggtgtcGAAaaagagacgtcgtcc | This work |
| *avrRpt2* D216E | Reverse | oCM35 | gacgtctctttTTCgacaccagtgaggacc | This work |

**Table S4** Summary of the type III effectors present in the *P. syringae* strains NY15125, DC3000, T1, NYT1, CA-407, and CA-A9.

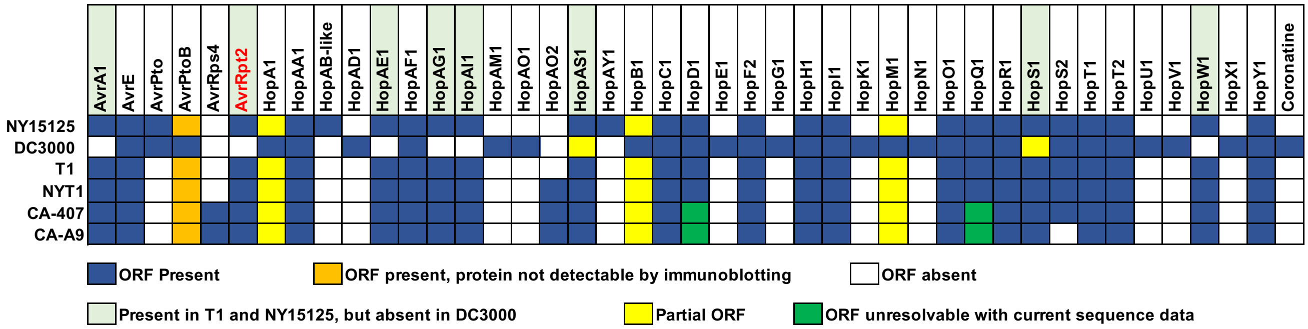


The type III effector genes are listed across the top and the *P. syringae* strains are listed at the left edge. A blue box indicates presence of a full length open-reading frame (ORF). A orange box indicates the gene is present but protein is not detectable by immunoblotting. A yellow box indicates the gene has an ORF truncated by an insertion sequence element or the presence of a premature stop codon. A green box indicates the ORF is unresolvable with current sequence data. A white box indicates the gene is not present in that strain. The light green shading indicates the effectors that are present in T1 (Almeida *et al.*, 2009), NYT1 (Jones *et al.*, 2015), NY15125 (Kraus *et al.*, 2017), CA-407, and CA-A9 (Kunkeaw *et al.*, 2010; Thapa *et al.*, 2015) but absent in DC3000 (Saha & Lindeberg, 2013) and were tested ion LA4245-R. AvrRpt2 is shown in red. DC3000 is the only strain that produces coronatine.

**Table S5** Features and expression patterns of the four genes in tomato that encode proteins with similarity to Arabidopsis RIN4.

| Gene | Solyc Identifier | No. amino acids | Predicted protein mass (kD) | % similarity to *At* RIN4 | PTI | | ratio | p value (FDR) | NTI | | ratio | p value (FDR) |
|---|---|---|---|---|---|---|---|---|---|---|---|---|
| | | | | | flgII-28 6 h | mock 6 h | | | RG-PtoR DC3000 6 h | RG-prf3 DC3000 6 h | | |
| Rin4-1 | Solyc09g059430 | 254 | 28.5 | 64.1 | 31.3 | 5.5 | 5.7 | 6.73E-19 | 25 | 14.7 | 1.7 | 0.0015 |
| Rin4-2 | Solyc06g083390 | 243 | 26.8 | 60.1 | 95.9 | 34.6 | 2.77 | 4.25E-08 | 90.5 | 27.9 | 3.2 | 6.68E-17 |
| Rin4-3 | Solyc12g098440 | 313 | 34.4 | 50.2 | 49.5 | 46.5 | 1.06 | 1 | 45.8 | 55.6 | 0.8 | 0.8179 |
| Rin4-4 | Solyc11g012010 | 224 | 24.8 | 59.7 | 0 | 0 | NA | NA | 0 | 0 | NA | NA |

The predicted protein mass is shown along with the percentage similarity to the Arabidopsis RIN4 protein (*At*RIN4). RNA-Seq data for PTI (PRR-triggered immunity) are from (Pombo *et al.*, 2014) and for NLR-triggered immunity (NTI) are from (Rosli *et al.*, 2013). Numbers under PTI and NTI are RPKMs (reads per kilobase of transcript per million mapped reads). FDR, false discovery rate. Data are available on the Tomato Functional Genomics Database (TGFD; http://ted.bti.cornell.edu/cgi-bin/TFGD/digital/home.cgi)

**Table S6** Gene models with the highest similarity to RPS2 and Mr5 in *S. lycopersicoides*.

(a)

| Gene Model | % similarity RPS2 | No. amino acid | Annotation | |
|---|---|---|---|---|
| Solyd08g052660 | 56.7 | 1,314 | NB-ARC | LRR |
| Solyd11g052900 | 56.1 | 743 | NB-ARC | LRR |
| Solyd10g073720 | 56.1 | 996 | NB-ARC | LRR |
| Solyd06g059160 | 54.8 | 740 | NB-ARC | LRR |
| Solyd02g054510 | 54.6 | 1027 | NB-ARC | LRR |

(b)

| Gene Model | % similarity Mr5 | No. amino acid | Annotation | |
|---|---|---|---|---|
| Solyd11g070490 | 69.3 | 916 | NB-ARC | LRR |
| Solyd11g069520 | 62.5 | 1329 | NB-ARC | LRR |
| Solyd03g059570 | 62.3 | 1226 | NB-ARC | LRR |
| Solyd11g069260 | 62.3 | 1431 | NB-ARC | LRR |
| Solyd11g070530 | 61.9 | 1244 | NB-ARC | LRR |

A genome-wide comparison between all proteins from *S. lycopersicoides* LA2951 and the amino acid sequences from RPS2 and Mr5 was performed to identify the most similar proteins present in this tomato accession. A pairwise alignment between each gene model and RPS2 (Kunkel *et al.*, 1993) (a) or Mr5 (Fahrentrapp *et al.*, 2012) (b) amino acids sequences was done to calculate the percentage of similarity.

**Supplemental Information References**


**Almeida NF, Yan S, Lindeberg M, Studholme DJ, Schneider DJ, Condon B, Liu H, Viana CJ, Warren A, Evans C, et al. 2009.** A draft genome sequence of *Pseudomonas syringae* pv. tomato T1 reveals a type III effector repertoire significantly divergent from that of *Pseudomonas syringae* pv. tomato DC3000. *Mol Plant-Microbe Interact* **22**(1): 52-62.

**Buell C, Joardar V, Lindeberg M, Selengut J, Paulsen I, Gwinn M, Dodson R, Deboy R, Durkin A, Kolonay J, et al. 2003.** The complete genome sequence of the Arabidopsis and tomato pathogen *Pseudomonas syringae* pv. tomato DC3000. *Proc Natl Acad Sci USA* **100**(18): 10181-10186.

**Fahrentrapp J, Broggini GAL, Kellerhals M, Peil A, Richter K, Zini E, Gessler C. 2012.** A candidate gene for fire blight resistance in Malus × robusta 5 is coding for a CC–NBS–LRR. *Tree Genetics & Genomics* **9**: 237-251.

**Jones LA, Saha S, Collmer A, Smart CD, Lindeberg M. 2015.** Genome-assisted development of a diagnostic protocol for distinguishing high virulence *Pseudomonas syringae* pv. tomato strains. *Plant Disease* **99**: 527-534.

**Kraus CM, Mazo-Molina C, Smart CD, Martin GB. 2017.** *Pseudomonas syringae* pv. tomato strains from New York exhibit virulence attributes intermediate between typical race 0 and race 1 strains. *Plant Disease* **101**: 1442-1448.

**Kunkeaw S, Tan S, Coaker G. 2010.** Molecular and evolutionary analyses of *Pseudomonas syringae* pv. tomato race 1. *Mol Plant-Microbe Interact* **23**: 415-424.

**Kunkel BN, Bent AF, Dahlbeck D, Innes RW, Staskawicz BJ. 1993.** RPS2, an Arabidopsis disease resistance locus specifying recognition of Pseudomonas syringae strains expressing the avirulence gene avrRpt2. *Plant Cell* **5**(8): 865-875.

**Kvitko BH, Collmer A. 2011.** Construction of *Pseudomonas syringae* pv. tomato DC3000 mutant and polymutant strains. *Methods Mol Biol* **712**: 109-128.

**Mahbou Somo Toukam G, Cellier G, Wicker E, Guilbaud C, Kahane R, Allen C, Prior P. 2009.** Broad diversity of *Ralstonia solanacearum* strains in Cameroon. *Plant Disease* **93**: 1123-1130.

**Nakagawa T, Suzuki T, Murata S, Nakamura S, Hino T, Maeo K, Tabata R, Kawai T, Tanaka K, Niwa Y, et al. 2007.** Improved Gateway binary vectors: high-performance vectors for creation of fusion constructs in transgenic analysis of plants. *Biosci Biotechnol Biochem* **71**(8): 2095-2100.

**Oh HS, Kvitko BH, Morello JE, Collmer A. 2007.** *Pseudomonas syringae* lytic transglycosylases coregulated with the type III secretion system contribute to the translocation of effector proteins into plant cells. *J Bacteriol* **189**: 8277-8289.

**Pombo MA, Zheng Y, Fernandez-Pozo N, Dunham DM, Fei Z, Martin GB. 2014.** Transcriptomic analysis reveals tomato genes whose expression is induced specifically during effector-triggered immunity and identifies the Epk1 protein kinase which is required for the host response to three bacterial effector proteins. *Genome Biol* **15**(10): 492.

**Rosli HG, Zheng Y, Pombo MA, Zhong S, Bombarely A, Fei Z, Collmer A, Martin GB. 2013.** Transcriptomics-based screen for genes induced by flagellin and repressed by pathogen effectors identifies a cell wall-associated kinase involved in plant immunity. *Genome Biol* **14**(12): R139.

**Saha S, Lindeberg M. 2013.** Bound to succeed: transcription factor binding-site prediction and its contribution to understanding virulence and environmental adaptation in bacterial plant pathogens. *Mol Plant Microbe Interact* **26**(10): 1123-1130.

**Thapa SP, Miyao EM, David MR, Coaker G. 2015.** Identification of QTLs controlling resistance to *Pseudomonas syringae* pv. tomato race 1 strains from the wild tomato, *Solanum habrochaites* LA1777. *Theor Appl Genet* **128**: 681-692.

**Whalen MC, Innes RW, Bent AF, Staskawicz BJ. 1991.** Identification of *Pseudomonas syringae* pathogens of *Arabidopsis* and a bacterial locus determining avirulence on both Arabidopsis and soybean. *Plant Cell* **3**: 49-59.

**Wroblewski T, Tomczak A, Michelmore R. 2005.** Optimization of Agrobacterium-mediated transient assays of gene expression in lettuce, tomato and Arabidopsis. *Plant Biotechnol J* **3**(2): 259-273.
